# The role of heavy metals in the co-selection of plasmid-borne metal and antibiotic resistance genes from industrially contaminated sediments

**DOI:** 10.64898/2025.12.14.694177

**Authors:** Brodie F. Gillieatt, Meghann Thai, Amy K. Cain, Ruth N. Zadoks, Nicholas V. Coleman, Michael A. Kertesz

## Abstract

A comprehensive understanding of the sources and drivers of antimicrobial resistance is essential for effective antimicrobial stewardship. Co-selection is now recognised as a significant driver of antimicrobial resistance, with established links between heavy metal exposure and the presence of bacteria with antimicrobial resistance. The precise mechanisms that drive this process in the environment are co-resistance, cross-resistance, and co-regulation, but their respective impacts remain largely unexplored. Here, we investigated whether heavy metal contamination in freshwater sediments selects for bacteria harbouring plasmids that carry both metal and antibiotic resistance genes, or genes encoding cross-resistance mechanisms. A diverse set of plasmids was recovered from metal-impacted sites at Lake Macquarie (New South Wales, Australia), which carried resistance genes particularly to copper, zinc, cobalt, cadmium, and arsenic. Two-thirds of these plasmids also carried one or more antibiotic resistance genes, indicating co-selection through the co-resistance mechanism. Functional assessment confirmed that the multi-metal and polymyxin resistance plasmid genotype was linked to the corresponding bacterial phenotype. The metagenome of the metal-impacted sediments was also examined to explore evidence of co-selection, and a broad range of incomplete plasmid sequences containing homologues of both metal- and antibiotic-resistance genes was detected. This study demonstrates the important link between anthropogenic heavy metal contamination and potentially clinically relevant antibiotic resistance genes. It highlights the importance of approaching the management of antimicrobial resistance from a One Health perspective.

## Introduction

The current antimicrobial resistance (AMR) crisis is on track to cause 10 million deaths per annum by 2050 [1]. The increasing incidence of AMR in bacterial infection is largely due to the acquisition of antibiotic resistance genes (ARGs), with many clinically relevant ARGs originating in the environment [2, 3]. Horizontal gene transfer (HGT) has mobilised resistance genes into clinical settings where they are propagated amongst commensal and pathogenic bacteria because of selective pressures exerted by antimicrobial use [4, 5].

In natural environments such as freshwater sediments, antibiotics are produced as a means of microbial competition and cell-to-cell signaling [6], and selective concentrations are strongly localised around the producing organism [2]. Heavy metals, by contrast, provide a less localised and more dominant selective agent in these environments. Heavy metals (defined here as metal elements with atomic number >20, density >5 g/cm^3^, plus arsenic and tellurium) are released from a variety of anthropogenic or natural sources and can accumulate in waterways, sediments, and soils at concentrations many orders of magnitude greater than antibiotics [7].

Bacterial resistance to these heavy metals is of considerable interest, as exposure to these compounds can select not only for metal resistance but also for AMR through a process known as co-selection. Three main models have been postulated for co-selection [7, 8]. The co-resistance model describes the acquisition of metal resistance genes (MRGs) and ARGs in tandem, typically by being co-localised on the same mobile genetic element (MGE). Alternatively, the cross-resistance model arises when a single gene product confers resistance to both metals and antibiotics. The third model, co-regulation, occurs when MRGs and ARGs share a common regulatory pathway. Evidence supporting metal and antibiotic co-selection includes the frequent observation that metal-resistant bacteria are often simultaneously resistant to antibiotics [9, 10], many MRGs are co-located with ARGs, indicative of co-resistance [11, 12], and efflux, detoxification, and sequestration resistance strategies are common to both antibiotic and heavy metal resistance, which is consistent with cross-resistance [3, 8]. Heavy metals can select for clinically relevant AMR pathogens *in vitro* [13] or in livestock [14], however, the prevalence and role of heavy metal resistance as driver of AMR in environmental settings needs to be understood better to address the global AMR crisis.

The mechanisms of co-selection, particularly co-resistance, are highly relevant in the investigation of plasmid-borne resistance. The mechanics of plasmid mobility are well-understood; conjugative plasmids can self-mobilise to other bacteria, mobilisable plasmids require mobilisation genes encoded *in trans*, and non-mobilisable plasmids cannot undergo conjugative transfer [15, 16]. Most plasmids exist as multiple copies per cell, thereby increasing the copy number of the resistance gene and potentially amplifying the resistance phenotype [17]. HGT of resistance genes can also be mediated by transposons, bacteriophage, and integrative and conjugative elements. The detection of resistance genes on plasmids and other MGEs in purified bacterial isolates is now routine, using sequence homology matches to entries in resistance gene databases such as CARD [18], AMRFinder [19], and BacMet [20]. Analysis of these resistance mechanisms is much more challenging when applied to the entire microbial community in a natural environment, since only approximately 1% of bacterial species are culturable using non-specific growth media [21]. Metagenomic sequencing provides a convenient methodology for studying bacterial and gene diversity of communities, yet it is constrained in functional assessment and manipulation of the detected gene sequences because sequences are often not associated with a specific bacterial host.

The waterways and sediments surrounding coal-fired power stations and industrial smelter sites are consistently exposed to elevated levels of heavy metals but have negligible exposure to antibiotics [22, 23]. These sites therefore provide an ideal environment for studying metal-resistant bacteria and evaluating the mechanisms of co-selection in the resistance plasmids they harbour. In this study, we have investigated freshwater sediments from two watercourses adjoining a coal-ash settling basin and a lead-zinc smelter in the Lake Macquarie region of New South Wales, Australia. The physicochemical properties of these sediments were analysed, and bacteria resistant to zinc, copper, or cadmium were isolated. Many of these isolates’ resistance phenotypes were due to the carriage of resistance plasmids enriched with MRGs, specifically for zinc, copper, and cadmium. Despite the absence of known anthropogenic sources of antibiotic selection, such as wastewater from municipal, hospital, or farm sites, many of the plasmids in these isolates also harboured ARGs. The genomic context of the resistance genes was analysed to identify evidence of co-resistance, cross-resistance, or co-regulation, and their importance in phenotypic regulation was confirmed by plasmid curing experiments.

## Materials and Methods

### Sampling and chemical analysis of sediments

Sediment samples were collected on 21 May 2021 from a creek bed leading directly downhill from the now decommissioned Pasminco Lead-Zinc Smelter (Site P: 32°56’25" S, 151°37’45" E) and from Crooked Creek, which receives groundwater from Eraring Power Station Coal Ash Basin (Site E: 33°3’41" S, 151°32’52" E). Both industrial sites have contributed significantly to the release of heavy metal particulates into the surrounding waterways and sediments over the past century [24–26]. Control sediment samples were taken from Lords Creek (Site L: 32°59’17" S, 151°29’20" E), an uncontaminated site in the same region. Triplicate samples (1 m apart) were taken from the top 50 mm of sediment to obtain relatively aerobic communities. Bacteria were cultured immediately upon arrival at the laboratory 4–6 hours after collection, and the remaining sediment was stored at 4 °C.

Moisture content was determined by the difference between wet and dry weight (75 °C, 48 h). Ion content was determined by ion chromatography of sediment extracts in deionised water. Cations (Na^+^, K^+^, NH ^+^, Ca^2+^, Mg^2+^) and anions (F^-^, Cl^-^, Br^-^, NO ^-^, NO ^-^, SO ^2-^, PO ^3-^) were measured using a Shimadzu LC-20A1 high performance liquid chromatography system with a CDD-10A conductivity detector and column oven (40 °C). Anions were eluted in 1 mM NaHCO_3_ /3.5 mM Na_2_CO_3_ (flow rate 1 mL/min) on a Dionex IonPac AS11 column (4 x 250 mm) with an AG11 guard column (4 x 50 mm), using a Shimadzu ICDS-40A suppressor. Cations were eluted in 500 mM oxalic acid (flow rate 1 mL/min) on a Shimadzu Shim-pack C4 column (4.6 x 150 mm) with no suppressor. Analysis of heavy metals (As, Cd, Co, Cr, Cu, Hg, Mn, Ni, Pb, Zn), reference elements (Al, Mg), total organic carbon, pH, and Eh was performed by Sydney Analytical Laboratories. Metal contamination was evaluated by calculating the geo-accumulation index [27] and enrichment factor [27] for each metal, with Lords Creek as background value and aluminium and magnesium as reference elements. The toxicity of metals was evaluated using the Nemerow integrated risk index (NIRI) [28], applying the toxic response of each element according to Hakanson [29].

### Microbial diversity of sediment samples

Total sediment DNA was extracted using the DNeasy PowerSoil Pro Kit (Qiagen, Germany) according to the manufacturer’s protocol, using five cycles of bead beating at 3000 rpm for 30 s (MBB-16 Biospec bead beater). Microbial diversity was analysed by amplicon sequencing of the 16S rRNA gene with 341F and 806R primers (Illumina MiSeq, Australian Genome Research Facility, Australia), and processed using DADA2 [30] and phyloseq [31]. Reads were assigned to taxa using the Silva v138.1 database [32], and amplicon sequence variants that appeared fewer than three times across all sites were removed.

### Isolation and identification of metal-resistant bacteria

Bacteria from sediments were grown for two weeks at ambient temperature on R2A agar containing 100 µg/mL cycloheximide and either 2.5 mM zinc (ZnSO_4_), 2.5 mM copper (CuSO_4_), or 0.5 mM cadmium (Cd(CH_3_CO_2_)_2_). To ensure divalent cations did not form insoluble phosphate salts, the R2A medium was amended by replacing 0.3 g/L K_2_HPO_4_ with 10 mM 3-(*N*-morpholino)propanesulfonic acid. High molecular weight DNA was then extracted from three colonies of each morphotype from each combination of site/heavy metal treatment, using a cetyltrimethylammonium bromide method [33], modified with a pretreatment step (1 mg/mL lysozyme, 0.3 mg/mL RNase A, 37 °C, 30 min), and protease addition during cetyltrimethylammonium bromide incubation (0.5 mg/mL proteinase K).

Strains were dereplicated by the comparison of BOX PCR amplicon band profiles [34], and the 16S rRNA gene of representative isolates was then sequenced using 341F and 805R primers (Australian Genome Research Facility, Australia). Sequences were aligned using MUSCLE in MEGA11 [35], and a phylogenetic maximum likelihood tree was constructed using 100 bootstrap replications and deleting nucleotide positions where more than 90% of sequences had a gap. *Cyanobacterium stanieri* PCC 7202 was used as a rooted outgroup.

### Genome sequencing, assembly and plasmid annotation

Isolated DNA from a total of 64 isolated strains was combined into 12 pools (2–8 isolates per pool) and Oxford Nanopore sequencing was performed at 900 Mb depth on R10.4.1 flow cells with Guppy v6.5.7 basecalling (SeqCenter, USA). The pooling was designed to minimise the phylogenetic relatedness between the isolates in each pool and reduce ambiguity in sequence assembly. Returned FASTQ reads above 1 kb in length were retained using Filtlong v0.2.1 [36] and quality was assessed using Nanoplot v1.36.2 [37]. Contigs were assembled with Flye v2.9.3 using automatic minimum overlap between read, metagenomic assembly, and one polishing iteration [38]. The individual contigs in each pool were assigned to the isolates in the pool by aligning either the entire contig sequence or the 16S rRNA gene sequence (identified using Barrnap [39]) to type strain sequences in the NCBI nucleotide collection. To verify plasmid sequence assembly, *Escherichia coli* UB5201 (R388) was included in Pooled Sample 1, and the resulting R388 sequence was found to be complete, with 99.74% identity to the original sequence (GenBank: BR000038).

Plasmid contigs were retrieved as circularised sequences smaller than 2 Mb that did not contain rRNA genes and had sequencing coverage ≥5. The plasmids present in the individual strains included in each sequencing pool were confirmed using PCR with primers specific for each plasmid (Table S1). Plasmids were assigned to known plasmid families using PlasmidFinder [40] and MOBscan [41]. Mobilisation genes (*oriT*, type four coupling proteins (T4CPs), relaxases and type four secretion systems (T4SSs)) were identified using oriTfinder [42]. Open reading frames (ORFs) were predicted using Prokka v1.14.6 [43], and annotated by applying a BLASTP alignment threshold of amino acid identity >30%, similarity >50%, query coverage >50%, and *E*-value <1×10^-20^. Specific gene names were assigned if amino acid identity exceeded 70%. ARGs were annotated using the CARD [18] and AMRFinder database [19], and MRGs were annotated using the BacMet experimentally confirmed database [20]. The BacMet, CARD, and AMRFinder database outputs were manually refined to retain only resistance genes that provided an experimentally characterised phenotype, as well as the removal of endogenous genes and biocide resistance genes.

All ORFs on a resistance plasmid were annotated based on alignments to the UniProtKB/Swiss-Prot database or the non-redundant protein sequences database. Specific gene names were only assigned if identity and query coverage exceeded 70%, otherwise a more generic gene family name was assigned applying a threshold of identity >30%, query coverage >50%, and *E*-value <1×10^-20^. BLASTP sequences that included specific gene names superseded lower *E*-value sequences that did not have a specific gene name, provided that the thresholds were still satisfied. MGEs within the plasmid sequences were identified using ISFinder [44], applying an *E*-value <1×10^-20^. The resulting putative insertion sequences (IS) were investigated further by manual examination to identify the inverted repeats flanking the IS. Additional inverted repeats were revealed by the Inverted Repeat Finder using ‘basic parameters’ [45]. Prophage sequences were identified using PHASTEST [46].

### Antibiotic and metal susceptibility

Antibiotic susceptibility was determined in triplicate following the Calibrated Dichotomous Susceptibility disc diffusion method [47] modified by incubating strains for 24 to 48 hours on R2A medium at 30 °C. The antibiotic discs applied were: amikacin 30 µg, ampicillin 10 µg, cefotaxime 30 µg, ceftriaxone 30 µg, chloramphenicol 30 µg, ciprofloxacin 5 µg, doxycycline 30 µg, erythromycin 15 µg, gentamicin 10 µg, imipenem 10 µg, kanamycin 30 µg, nalidixic acid 30 µg, neomycin 30 µg, spectinomycin 25 µg, streptomycin 25 µg, compound sulfonamides 300 µg, tetracycline 30 µg, and trimethoprim 5 µg (ThermoFisher Scientific, USA). An inhibition annular radius of <6 mm was recorded as resistant, and ≥6 mm as susceptible, as recommended [47].

Strains that presented a multi-antibiotic resistance phenotype were cured of their plasmids by exposure to acridine orange (2.5–640 µg/mL). Cultures were incubated in the dark for two days at 5–10 °C above their optimum growth temperature and then plated onto LB or R2A agar with or without selective agents (2.5 mM zinc, 2.5 mM copper, or 0.5 mM cadmium). Loss of the plasmid in any resulting metal-sensitive colonies was confirmed by PCR using plasmid-specific primers (Table S1).

Minimum inhibitory concentrations (MICs) were measured according to the EUCAST guidelines [48], modified to accommodate optimal growth. Mid-exponential growth phase cultures were inoculated (OD_600_ = 0.01) into 500 µL of growth medium in a 48-well polystyrene microtitre plate (*Sphingomonas hankookensis* – 5-fold diluted LB medium; *Pigmentiphaga litoralis* –amended tryptic soy broth medium (20 g/L bacteriological peptone, 2.2 g/L glucose, 5 g/L NaCl, 10 mM 3-(*N*-morpholino)propanesulfonic acid, pH 7)). The following metals or antibiotics were added at appropriate serial dilutions: zinc (ZnSO_4_), copper (CuSO_4_), cadmium (Cd(CH_3_CO_2_)_2_), cobalt (CoCl_2_), nickel (NiCl_2_), arsenate (Na_3_AsO_4_), arsenite (NaAsO_2_), mercury (HgCl_2_), ampicillin sodium salt, polymyxin B sulfate (both antibiotics prepared daily). The cultures were incubated at 30 °C, 200 rpm for 18 h for *S. hankookensis*, or 48 h for *P. litoralis*, and growth was measured as OD_600_ with a Clariostar Plus plate reader (BMG Labtech, Germany). The MIC was defined as the antimicrobial concentration sufficient to inhibit growth to below OD_600_ = 0.1 at the end of the growth period.

Functional β-lactamase activity in whole cells was tested using nitrocefin [49], using *E. coli* HB101 (pET15bVP) as a positive control and *E. coli* S17-1 as a negative control. Where required, strains were pre-exposed to ampicillin by growing them on an LB plate for 48 hours in the presence of an ampicillin disc (10 μg) to induce β-lactamase activity, and then testing bacteria growing adjacent to the disc for β-lactamase activity.

### Cultivation-independent plasmid extraction and characterisation from sediments

Sediment (30 g) from each site was spiked with 2 or 20 mM cadmium (Cd(CH_3_CO_2_)_2_) or chromium (K_2_Cr_2_O_7_), and incubated at 25 °C for 28 days, maintaining moisture content by weekly additions of water as required. Plasmid DNA was then extracted from the bacterial fraction of both treated and original sediments, as described below. A plasmid-free extraction control was also included (*E. coli* TOP10 inoculated into autoclaved sterile sand and allowed to adhere for 3 h before extraction).

Extraction of plasmids from sediment and sand was modified from Heringa et al [50]. Bacteria were extracted by homogenising sediment with 5 volumes of phosphate buffered saline (PBS) containing 0.5% v/v Tween-20, pH 8.5 for 3 x 1 min cycles with 1 min rest intervals at 4 °C (500 W Anko BL9706-CB blender). The resulting slurry was layered onto an equal volume of Nycodenz solution (ProteoGenix, France, 800 g/L in PBS, 0.3 mM EDTA) and centrifuged (14,000 *g*, 30 min, 4 °C). The interface layer was removed, mixed with two volumes of PBS and centrifuged again (17,000 *g*, 20 min). Total DNA was extracted from the resulting bacterial pellet by 1 h incubation at 37 °C in 500 µL PBS supplemented with lysozyme (1 mg/mL) and RNase A (0.3 mg/mL), followed by phenol/chloroform extraction and ethanol precipitation. Chromosomal DNA was removed by incubation (37 °C, 3 h) with 10 U of plasmid-safe-ATP-dependent DNase (Astral Scientific, Australia), and the remaining plasmid DNA was amplified using the REPLI-g Mini Kit (Qiagen, Germany), following the manufacturer’s protocol.

The amplified DNA was subjected to Oxford Nanopore sequencing to a depth of 300 Mb (SeqCenter, USA). The resulting sequences were assembled into contigs as described above, with additional removal of reads in the bottom 10% of read quality using Filtlong. Contigs that aligned with the no plasmid control (identity >70%, query coverage >50%, *E*-value <0.001) were manually removed from the contig collection, along with contigs containing rRNA genes or with <5-fold coverage [51]. ORFs were annotated as described above.

### Data processing

All data processing was performed in R 4.5.2, unless indicated otherwise, with a significance threshold of *p* <0.05 for all statistical tests. Normality of data was assessed with a Shapiro-Wilk test. Equality of variance was assessed by Levene’s test. Significance of normal data with equal variance was determined by an ANOVA with Tukey’s post hoc test.

Significance of non-normal data was determined by a Kruskal Wallis test and Dunn’s post hoc test with Benjamin-Hochberg *p-*value correction. Variable relationships were evaluated by Spearman’s coefficient and redundancy analysis using vegan package version 2.6-4 [52] with 10,000 ANOVA permutations. Alpha diversity was calculated using Shannon’s and Simpson’s indices with 95% confidence intervals calculated using Hutcheson’s t-test. All plots were visualised using the ggplot2 package [53]. Plasmid sequences were visualised on SnapGene 8.0.2.

## Results

### Industrial sediments showed elevated heavy metal loads and greater numbers of cultivable metal-resistant bacteria

The mean total heavy metal load of the sediments was higher at the two industrial sites than at Site L (Fig. S1), but no individual heavy metal was significantly elevated at both industrial sites. The geo-accumulation indices classified both industrial sites as slightly to moderately polluted with arsenic, chromium, and mercury, and Site P sediment was also slightly polluted with cadmium and lead (Table 1). The mercury levels at Site P were attributed to anthropogenic contamination according to the enrichment factor, and mercury toxicity posed an ecological risk at both industrial sites (NIRI [29]), as did the cadmium level at Site P.

**Table 1:** Heavy metal contamination in industrial sediments indicates moderate pollution and ecological risk. Geo-accumulation index (I_geo_) [27], enrichment factor (EF) [27] and Nemerow integrated risk index (NIRI) [28] were calculated based on measured metal concentration in sediment samples. Sediment from Site L was used as the background reference.

| Index | Site P | Site E |
| --- | --- | --- |
| $I_{geo}$ | As, Cd, Cr, **Hg**, Pb | As, **Cr**, Hg |
| EF | Hg |  |
| NIRI | Cd, **Hg** | Hg |

Element – Slightly polluted ( $I_{geo}$ 0–1)/minor enrichment (EF 1–3)/ moderate risk (NIRI 40–80)
**Element** – Moderately polluted ( $I_{geo}$ 1–3)/moderate enrichment (EF 3–5)/considerable risk (NIRI 80–160)
**Element** – Strongly polluted ( $I_{geo}$ 3–5)/moderately severe enrichment (EF 5–10)/high risk (NIRI 160–320)
**Element** – Extremely polluted ( $I_{geo}$ >5)/severe enrichment (EF >10)/extreme risk (NIRI >320)

Culturable, cadmium- or copper-resistant bacteria were more abundant in the industrial sediments than at Site L, both in terms of absolute and relative colony forming units (CFUs) (Fig. 1). In contrast, zinc-resistant bacteria were similarly abundant at all three sites. The mercury, cadmium, and lead contents of each sediment sample were positively correlated (Spearman coefficient >0.4) with the numbers of cadmium-resistant and copper-resistant CFUs (Fig. S2), though this correlation was only statistically significant for mercury (*p* = 0.00001). Chromium, mercury, and arsenic contents were positively associated with the number of zinc-resistant CFUs, although only the correlation with chromium was statistically significant (*p* = 0.033). Interestingly, zinc, cobalt, and manganese concentrations were negatively correlated (Spearman coefficient <-0.4) with metal-resistant CFUs, while copper and nickel contents displayed no notable correlations.

**Fig. 1:**
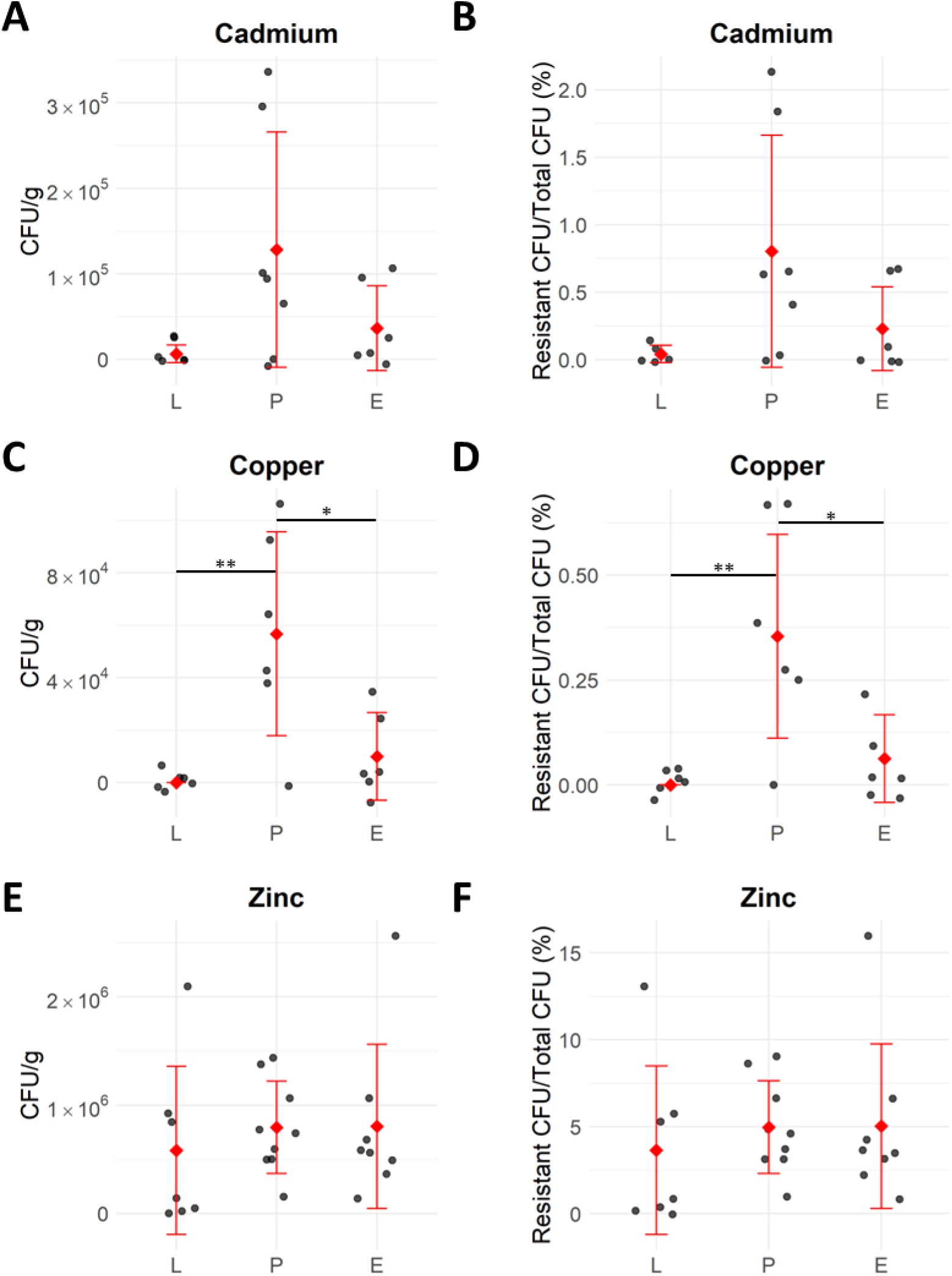
Cultivable metal-resistant bacteria in freshwater sediments from industrial (P, E) and control (L) sites. Colony-forming units (CFU) were determined on R2A agar supplemented with either (**A**, **B**) 0.5 mM cadmium acetate, (**C**, **D**) 2.5 mM copper(II) sulfate, or (**E**, **F**) 2.5 mM zinc sulfate. Left-hand panels show total populations (CFU/g), right-hand panels show the relative populations. Black circles represent individual replicates (*n* = 6–8), red diamonds represent the mean with standard deviation error bars (*p* values: **<0.01, *<0.05, Kruskal Wallis test).

A bacterial collection of 88 unique strains was compiled from these metal-resistant isolates. All but one strain could be identified by comparison of the V3-V4 region of the 16S rRNA to type strain sequences to genus level (>97% sequence identity), and two-thirds of the strains could be identified to species level (>99% sequence identity) (Table S2). These strains belonged to one of four phyla: *Actinomycetota*, *Pseudomonadota*, *Bacteroidota*, and *Bacillota* (Fig. 2). *Actinomycetota* was primarily represented by *Streptomycetales* and *Micrococcales*, while *Pseudomonadota* was mainly represented by *Hyphomicrobiales* and *Burkholderiales*. *Bradyrhizobium*, *Cupriavidus*, *Methylobacterium*, *Mycolicibacterium*, and *Streptomyces* were found in both industrial sediments, suggesting these are common, metal-resistant, freshwater sediment-dwelling microbes. Site P sediment contained the highest diversity of metal-resistant, culturable bacteria, with nine orders and twenty-one genera. Site E and the uncontaminated Site L displayed less diversity with six orders and ten genera, or five orders and nine genera, respectively.

**Fig. 2:**
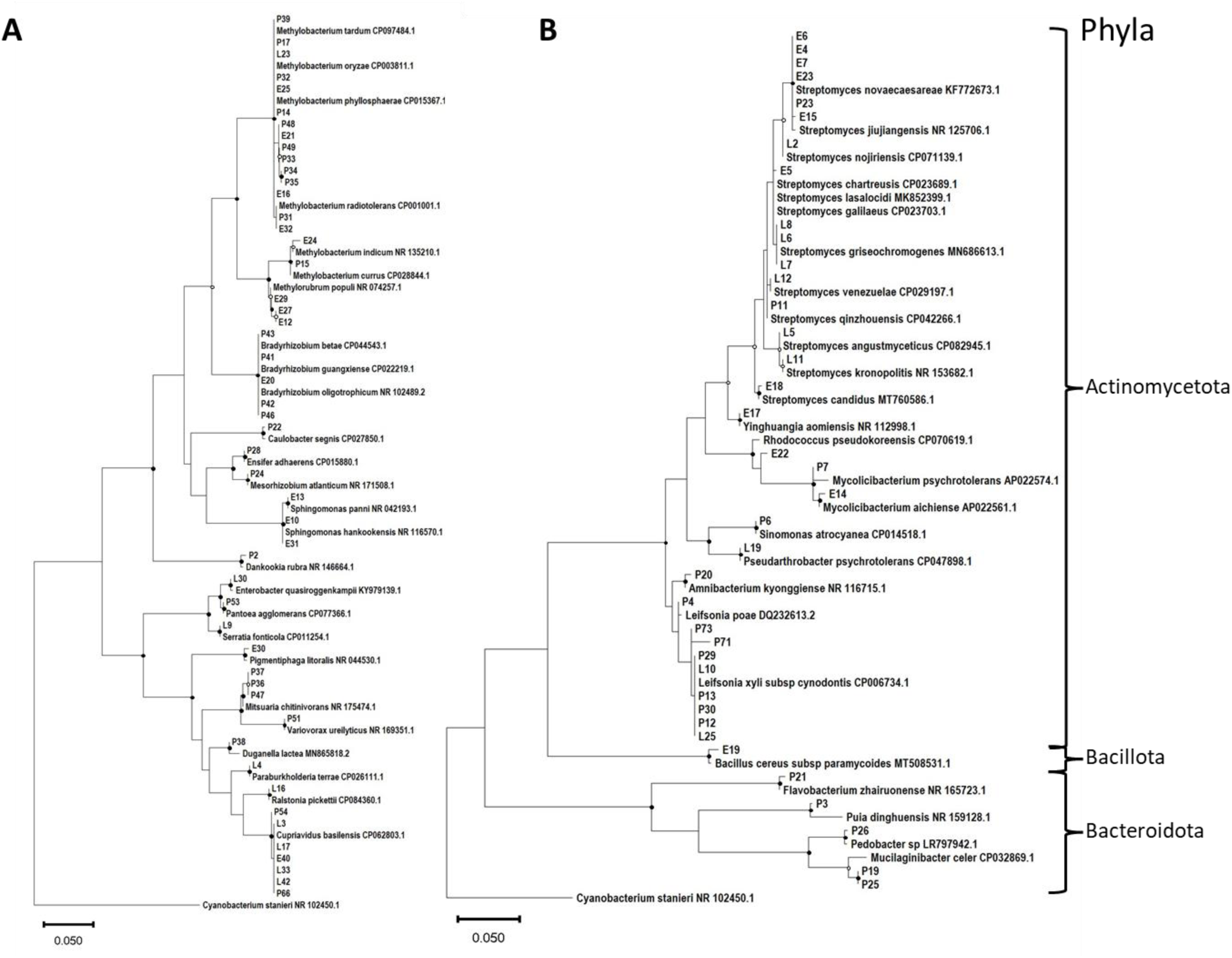
Maximum likelihood phylogenetic trees of unique metal-resistant bacterial isolates. Isolates obtained in this study are labelled by site (L, P, E) and sequential isolate number. 16S rRNA gene of isolates is aligned with the closest matching type strain from the National Center of Biotechnology Information. Isolates of the (**A**) *Pseudomonadota* phylum and (**B**) other phyla are displayed in separate trees, both rooted to *Cyanobacterium stanieri* PCC 7202. ○ >70% bootstrap support, ● >85% bootstrap support.

The culturable metal-resistant bacteria present in the sediments only represented a subset of the total bacterial diversity present. This was measured using DNA extracted directly from the sediment, and the total bacterial population was found to be dominated by common soil dwelling bacteria at all sites, with an imbalance in the dominance of *Arthrobacter* present in the industrial sediments than at Site L (Fig. S3). The alpha diversity measures reflected this, with higher values at Site L than at the industrial sites (Fig. S4), which contrasts to the diversity of cultured metal-resistant isolates.

### Strains from industrial sediments contained a range of large, novel, metal-resistance plasmids

Whole genome sequences were obtained for 64 isolates, representing every genus in the collection, and yielded an average of 35.5 contigs per isolate (Table S3). Fifty-nine complete plasmid sequences were identified, ranging in size from 4.3 kb to 1.9 Mb, with a 90% mean of 106 kb (Fig. S5). Plasmid sizes varied by taxonomic order, with the three most common orders (*Hyphomicrobiales*, *Sphingomonadales*, and *Micrococcales*) containing large plasmids (>50 kb). At least 28 plasmids were obtained from Site P, 24 from Site E, and 5 from Site L (Table S4). Three pairs of identical plasmids (>99% identity and query coverage) were found in more than one strain; *Sphingomonas* isolates E10 and E31 shared two plasmids, and isolates E10 and E13 shared one plasmid.

No REP or Inc types could be determined for these plasmids, so they were classified by MOB typing, which is based on the relaxase gene [41]. The plasmids predominantly belonged to the MOB_F_ or MOB_P_ types, although six contained relaxase genes that are not associated with any known MOB-type, and twenty-five lacked a relaxase gene (Table S4). Plasmid types were not exclusively associated with any taxonomic order or site (Fig. S6).

Alignment of plasmid ORFs with resistance gene databases revealed that approximately one-third of the plasmids carried putative MRGs or ARGs (Table 2). Ten resistance plasmids were from Site P and eleven were from Site E, and all were novel, with the greatest overlap to a known plasmid being 54% (Table S4). No resistance plasmids were retrieved from Site L. Of total ORFs, 13.2% were MRGs and most plasmids carried multiple resistance gene types. The most frequently occurring MRGs conferred multi-metal, copper, and arsenic resistance (Fig. 3A), and were present on 80%, 60%, and 45% of plasmids, respectively (Fig. 3B). Copper resistance was mainly represented by *cop* and *cus* operons, while multi-metal resistance was dominated by the *czc* operon, which encodes resistance to cadmium, zinc and cobalt [54]. Combining the abundance of *czc* and zinc-specific resistance genes, the number of zinc resistance genes was equivalent to copper resistance genes.

**Fig. 3:**
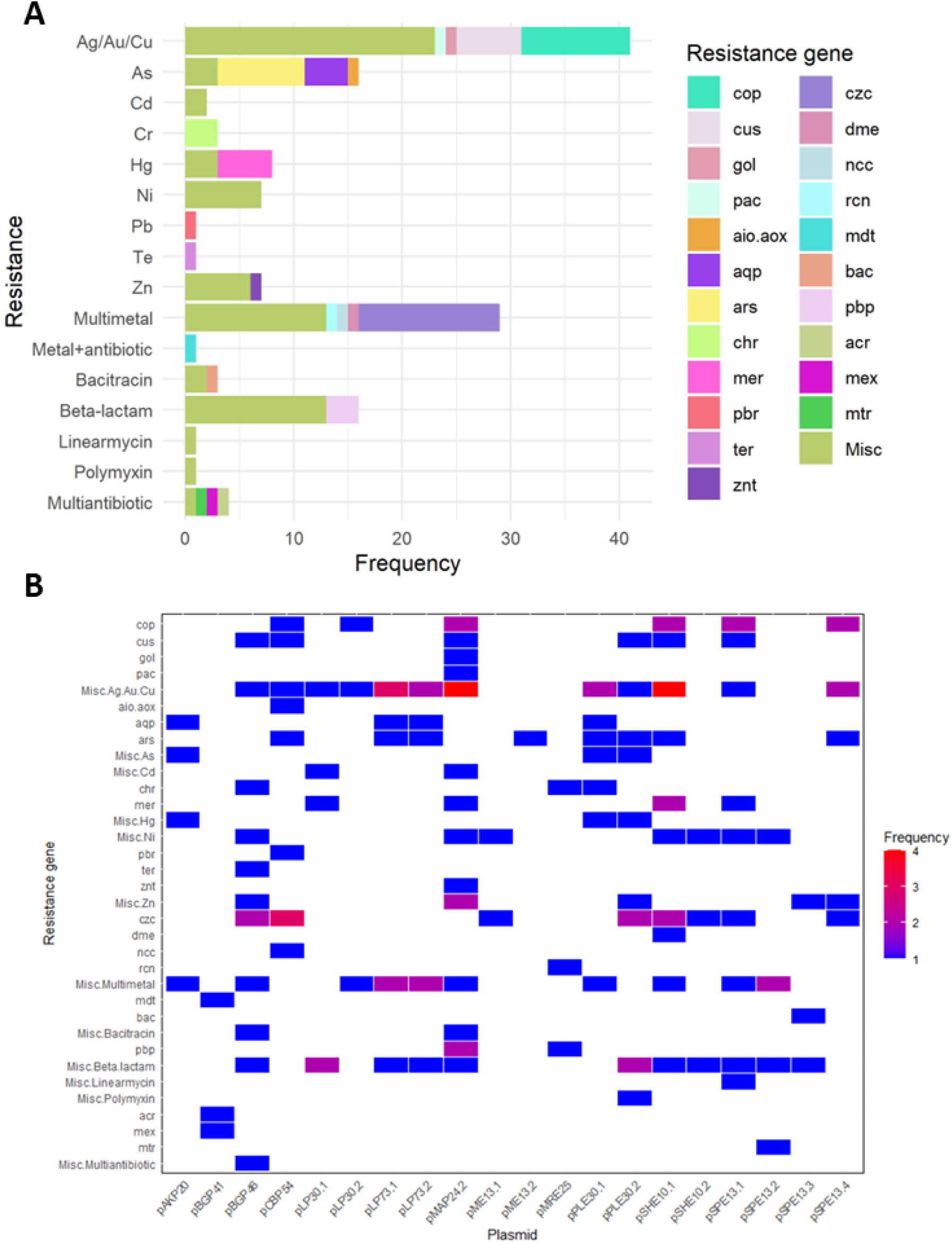
Copper, arsenic, multi-metal, and β-lactam resistance genes predominated in plasmids from metal-resistant isolates. (**A**) Resistance genes identified on plasmids classified by their predicted target substrate. (**B**) Heat map displaying the distribution and copy number of resistance genes per plasmid. Plasmid pPAP53 was excluded due to incomplete annotation, attributed to its large size. ‘Misc’ = miscellaneous resistance genes with insufficient identity to any specific, characterised resistance gene.

**Table 2:** Features of resistance plasmids recovered from metal-resistant isolates. Plasmid contig size, mobility (MOB) type, Oxford Nanopore sequencing coverage, host identification by PCR, and annotated resistance and mobilisation genes are displayed. MOB type was predicted by MOBscan [41], based on relaxase gene identity; plasmids without relaxase genes are classified as not applicable (NA), whereas plasmids with relaxase genes that did not belong to a known group are classified as novel. Metal resistance genes (MRG), and antibiotic resistance genes (ARG) were identified from database alignments using BacMet [20], CARD [18], and AMRFinder [19]. Miscellaneous (misc) genes have a lower alignment score than specifically named resistance genes. MBL = metallo-β-lactamase, T4CP = type four coupling protein, T4SS = type four secretion system

| Plasmid | Size (bp) | MOB type | Sequence coverage | Host | MRG families | ARG families | Mobility elements | GenBank accession number |
| --- | --- | --- | --- | --- | --- | --- | --- | --- |
| pCP54 | 171,550 | NA | 7 | *Cupriavidus* sp. P54 | *aio/aox, ars, cop, cus, czc, ncc, pbr*, misc. Cu resistance genes |  |  | PQ035965 |
| pAKP20 | 51,845 | P | 144 | *Amnibacterium kyonggiense* P20 | *aqp*, mics. As, Hg and multi-metal resistance genes |  | Relaxase gene | OP178894 |
| pMRE25 | 46,586 | F | 11 | *Methylobacterium radiotolerans* E25 | *chr, rcn* | PBP gene | Relaxase, T4CP genes | OQ588744 |
| pBGP41 | 248,638 | P | 25 | *Bradyrhizobium guangxiense* P41 | *mdt* | *acr, mdt, mex* | Relaxase, T4CP, complete T4SS genes | PQ035969 |
| pPLE30.1 | 100,066 | P | 52 | *Pigmentiphaga litoralis* E30 | *aqp, ars, chr*, misc. As Cu, Hg, resistance genes |  | Relaxase gene | PQ035964 |
| pPLE30.2 | 199,488 | NA | 57 | *P. litoralis* E30 | *ars, czc, cus*, misc. As, Cu, Hg, Zn resistance genes | *bla*<sub>OXA-1383</sub>, *mcr-12*<sup>+</sup> |  | PQ035968 |
| pLP30.1 | 57,599 | P | 92 | *Leifsonia* sp. P30 | *mer*, misc. Cd, Cu resistance genes | MBL gene | Relaxase, partial T4SS genes | PQ035966 |
| pLP30.2 | 128,795 | NA | 105 | *Leifsonia* sp. P30 | *cop*, misc. Cu, multi-metal resistance genes | *mdt*-homologue | Relaxase, T4CP, partial T4SS genes | PQ035967 |
| pMAP24.2 | 381,974 | Novel | 8 | *Mesorhizobium atlanticum* P24 | *cop, cus, gol, mer, pac, znt*, misc. Cd, Cu, Ni, Zn resistance genes | *bac*-homologue, MBL, PBP genes | Relaxase, T4CP, partial T4SS genes | PQ196794 |
| pBGP46 | 180,240 | P | 43 | *B. guangxiense* P46 | *chr, cus, czc, ter*, misc. Ag, Cu, Ni, Zn, multidrug resistance genes | *acr*-homologue, *bac*-homologue, MBL genes | Relaxase, T4CP, complete T4SS genes | PQ196795 |
| pSHE10.1 | 202,926 | F | 149 | *Sphingomonas hankookensis* E10, E31 | *ars, cop, cus, czc, dme, mer*, misc. Ag, Cu, Ni, Zn resistance genes | MBL gene | Relaxase, T4CP genes | PQ196796 |
| pSHE10.2 | 64,252 | P | 173 | *S. hankookensis* E10 | *czc*, misc. Ni resistance genes | MBL gene | Relaxase, T4CP, complete T4SS genes | PQ196797 |
| pPAP53 | 1,892,539 | NA | 44 | *Pantoea agglomerans* P53 | *act, ars, cop*, misc. Ag resistance genes |  | T4CP gene | Not submitted, incomplete annotation |
| pSPE13.1 | 206,010 | NA | 294 | *Sphingomonas panni* E13 | *cop, cus, czc, mer*, misc. Cu, Ni, multi-metal resistance genes | Linearmycin resistance, MBL genes |  | PQ201170 |
| pSPE13.2 | 69,420 | F | 531 | *S. panni* E13 | Misc. Ni, multi-metal resistance genes | *mtr*, MBL gene | Relaxase gene | PQ201171 |
| pSPE13.3 | 38,227 | NA | 348 | *S. panni* E13 | Misc. Zn resistance gene | *bac*, misc. β-lactamase gene | T4CP gene | PQ201172 |
| pSPE13.4 | 241,771 | F | 213 | *S. panni* E13 | *ars, cop, czc*, misc. Cu, Zn resistance genes |  | Relaxase, T4CP genes | PQ201173 |
| pME13.1 | 64,947 | Novel | 14 | *Methylobacterium* sp. | *czc*, misc. Ni resistance gene |  | Relaxase, T4CP, complete T4SS genes | Not submitted, unknown host |
| pME13.2 | 48,756 | F | 7 | *Methylobacterium* sp. | *ars* |  | Relaxase gene | Not submitted, unknown host |
| pLP73.1 | 68,729 | P | 19 | *Leifsonia* sp. P73 | *aqp, ars*, misc. Cu, multi-metal resistance genes | *cmcT*-homologue | Relaxase gene | PQ201174 |
| pLP73.2 | 69,333 | P | 15 | *Leifsonia* sp. P73 | *aqp, ars*, misc. Cu, multi-metal resistance genes | *cmcT*-homologue | Relaxase gene | PQ201175 |

ARGs were detected on two-thirds of the plasmids but constituted only 1.1% of total ORFS. Most detected ARGs were β-lactamase genes or multi-antibiotic efflux pump genes, which were found on 55% and 15% of plasmids, respectively. A single gene encoding a cross-resistance mechanism was identified (*mdtA* on pBGP41), which is thought to confer resistance to copper, zinc, novobiocin, and β-lactams [55, 56].

The antibiotic susceptibility of strains carrying resistance plasmids was tested with 18 different antibiotics from 7 classes (Table S5). The most common phenotype was nalidixic acid resistance, with seven strains displaying resistance (as defined in the Calibrated Dichotomous Susceptibility methods, not in the clinical sense). *Bradyrhizobium guangxiense* P46 exhibited resistance to the most antibiotics, with resistance to seven antibiotics from six classes. *Methylobacterium radiotolerans* E25 and *Cupriavidus basilensis* P54 were resistant to five antibiotics from four classes, and four antibiotics from four classes, respectively. Interestingly, although β-lactamase genes were the most common predicted ARG carried on these plasmids, none of the strains expressed a functional β-lactamase even after pre-exposure to ampicillin, as tested by nitrocefin hydrolysis.

In order to evaluate whether the resistance phenotypes correlated with the resistance plasmid genotype, strains that were resistant to at least two antibiotics were subjected to plasmid curing and then susceptibility re-testing. Plasmid removal was only successful for *P. litoralis* E30, which lost pPLE30.2 but not pPLE30.1, and *S. hankookensis* E10, which lost both pSHE10.1 and pSHE10.2. The pPLE30.2 plasmid contained predicted MRGs for cadmium, zinc, cobalt, copper, and arsenic, as well as the novel resistance genes, *mcr-12* and *bla*_OXA-1383_. Both *mcr-12* and *bla*_OXA-1383_ are characterised in Gillieatt et al [57]. The polymyxin B MIC decreased 32-fold for the plasmid-cured strain, confirming the polymyxin resistance function of *mcr-12* (Table 3, Fig. S7A; from Gillieatt et al [57]). The plasmid-cured *P. litoralis* exhibited a 32-fold reduction in MIC for arsenate and arsenite for the plasmid-cured *P. litoralis*, highlighting the significant role of *arsHCB* in mediating arsenic resistance (Fig. S7B–C). A two-, two-, and four-fold reduction in MIC compared to the wild-type for cadmium, cobalt, and zinc, respectively, was likely due to the loss of *czcCBA* and *czcD*-homologues (Fig. S7D–F). The MIC for copper was identical for both plasmid-cured and wild-type strains despite the loss of *cusAB* and *copABCDG*-homologue genes (Fig. S7G).

**Table 3:** Minimum inhibitory concentrations (MICs) of heavy metals and antibiotics for *Pigmentiphaga litoralis* and *Sphingomonas hankookensis* wild-type and plasmid-cured strains. MICs were determined based on OD_600_ <0.1 following 48 hours of growth in amended tryptic soy broth (*P. litoralis*), or 18 hours growth in 1/5 strength Luria-Bertani medium (*S. hankookensis*). *n* = 2*–*3.

| Antimicrobial | Wild-type strain<br>MIC (mM) | Plasmid-cured strain<br>MIC (mM) | Fold-change<br>between MICs | Likely plasmid<br>genes |
| --- | --- | --- | --- | --- |
| *Pigmentiphaga litoralis*<sup>†</sup> |  |  |  |  |
| Arsenate | 128 | 4 | -32 | *arsHCB* |
| Arsenite | 5 | 0.156 | -32 | *arsHCB* |
| Cadmium | 2 | 1 | -2 | *czcCBA* |
| Cobalt | 2 | 1 | -2 | *czcCBA* |
| Copper | 5 | 5 | No change | *cusAB, copABCDG* |
| Zinc | 8 | 2 | -4 | *czcCBA* |
| Polymyxin B | 80 µg/mL | 2.5 µg/mL | -32 | *mcr-12* |
| *Sphingomonas hankookensis* |  |  |  |  |
| Arsenate | 128 | 4 | -32 | *arsHNB* |
| Arsenite | 2 | 0.125 | -16 | *arsHNBC* |
| Cadmium | 0.125 | 0.031 | -4 | *czcCBAP* |
| Cobalt | 0.5 | 0.063 | -8 | *czcCBAP, dmeF* |
| Copper | 1.25 | 0.625 | -2 | *copDCBAG, cusAB* |
| Mercury | 0.015 | 0.008 | -2 | *merTPA* |
| Nickel | 1 | 0.25 | -4 | *dmeF* |
| Zinc | 2 | 0.5 | -4 | *czcCBAP* |
| Ampicillin | 25 µg/mL | 25 µg/mL | No change | Metallo-β-lactamase gene |

<sup>†</sup>MIC phenotypes reported in [57]

Similar results were seen for the plasmids harboured by *S. hankookensis* E10 (pSHE10.1 and pSHE10.2). These plasmids carried MRGs involved in resistance to arsenic, cadmium, cobalt, copper, mercury, nickel, and zinc, as well as β-lactamase genes. The *S. hankookensis* plasmid-cured strain displayed substantial MIC reductions for arsenate and arsenite (32-fold and 16-fold, respectively) likely due to the loss of *arsHNBC* (Table 3, Fig. S8A–B). A four-, eight-, and four-fold reduction in MIC compared to the wild-type for cadmium, cobalt, and zinc respectively was likely due to the loss of *czcCBAP* (Fig. S8C–E). Similarly, copper, mercury, and nickel MICs decreased by two-, two-, and four-fold, respectively, likely due to the loss of *copDCBAG*, *merTPA*, and *dmeF* respectively (Fig. S8F–H). The ampicillin MIC was identical for both strains (25 µg/mL).

For the remaining strains, the inability to obtain a metal-sensitive phenotype under plasmid-curing conditions indicates either that the resistance phenotype was not solely due to plasmid-borne resistance genes, or that the resistance plasmid was essential and could not be removed under the conditions used.

### Genetic recombination between resistance plasmids in sediments

In three cases, pairs of unique resistance plasmids from the same site and same host genus were found to share significant regions of homology, consistent with recombination having occurred between them. Two plasmids from a *Leifsonia* isolate, pLP73.1 and pLP73.2, shared 96.48% identity across 51.8 kb in 3 regions of the plasmids: a 16.3 kb resistance region containing copper resistance genes, arsenic resistance genes and a *cmcT*-homologue; a 22.1 kb T4SS and *rep* region; and a 13.4 kb region containing cation diffusion facilitator and relaxase genes, adjacent to a recombinase gene present on both plasmids. The homologous T4SS and *rep* region was bordered by a IS*30* transposase gene on pLP73.2, but not on pLP73.1, indicating that this may have been involved in a cut-and-paste transposition of this section but is no longer intact.

*Methylobacterium* plasmids pME13.2 and pMRE25, which are from different isolates, shared 81% identity across 10.9 kb. This was subdivided on the plasmids into a 6.8 kb region containing a relaxase and T4CP gene, and a 4.1 kb region containing *parA* and a plasmid retention toxin gene. The shared relaxase gene region was within 5 kb of an IS*Mex* family inverted repeat and transposase gene on pMR13.2, and an IS*Mtsp5* transposase gene on pMRE25, which may be involved in its transposition.

The final homologous plasmid pair, pSHE10.1 and pSPE13.1, originate from different *Sphingomonas* isolates, and share 99.78% identity across 76.2 kb. The largest homologous region on the plasmids was a 50.7 kb resistance region including *czc*, *cop*, and *cus* operons and a metallo β-lactamase gene. This sequence was flanked on both plasmids by a Tn*5393* transposase gene on one end and an IS*110* or IS*1182* family transposase gene on the other end. These may be involved in copy-and-paste transposition of this fragment.

These three examples all reveal the presence of transposase genes in the homologous regions. Transposase-like genes were carried on most of the plasmids, but only eight plasmids carried a transposon or IS with complete detectable inverted repeats (Table S6). None of the resistance genes were obviously within transposons, although some transposons were adjacent to resistance genes. For example, a β-lactamase gene was adjacent to the left flank of Tn*5393* on pSHE10.1 (Fig. S9A), IS*MPo10* on pSPE13.2, and Tn*5393* on pSPE13.1. It is possible the β-lactamase gene is within a transposon which has lost an inverted repeat as only a right inverted repeat was detected. Similarly, the right inverted repeat of IS*Gbe1* on pSHE10.1 and pSPE13.4 was adjacent to a copper oxidase gene (Fig. S9B). Four potential novel IS were identified in *Sphingomonas panni* plasmids, marked by novel inverted repeats flanking a transposase gene. No bacteriophage specific genes were identified on the plasmids.

### Heavy metal and antibiotic co-resistance encoded on plasmids

Five of the plasmids studied were notable because they not only carried both MRGs and ARGs, indicative of co-resistance, but also had many hallmarks of mobilisable or conjugative plasmids. This is suggestive of the development of co-resistance in the industrial sediments with the risk of HGT to other bacterial hosts. Plasmids pBGP41, pBGP46, pSHE10.2, and pME13.1 contained all genetic elements for conjugation (relaxase, T4CP, T4SS), apart from an *oriT*, and a further 14 carries some of these elements indicating that they may be mobilisable (Table 2).

*Pigmentiphaga litoralis* contained two resistance plasmids as described above, with pPLE30.2 providing evidence of co-resistance. This 199.5 kb plasmid, of an unknown type, harbours a 50 kb region rich in MRGs demonstrated to provide functional resistance to not just metals, but polymyxin B (Table 3; also see [57]). Notably, *mcr-12* is directly flanked downstream by a *czcCBA*-homologue and upstream by a *cusRS*-like two-component regulator and *ars* operon, consistent with co-selection by co-resistance (Fig. 4A). It is possible that the *cusRS*-homologue regulates expression of all these genes consistent with co-regulation.

**Fig. 4:**
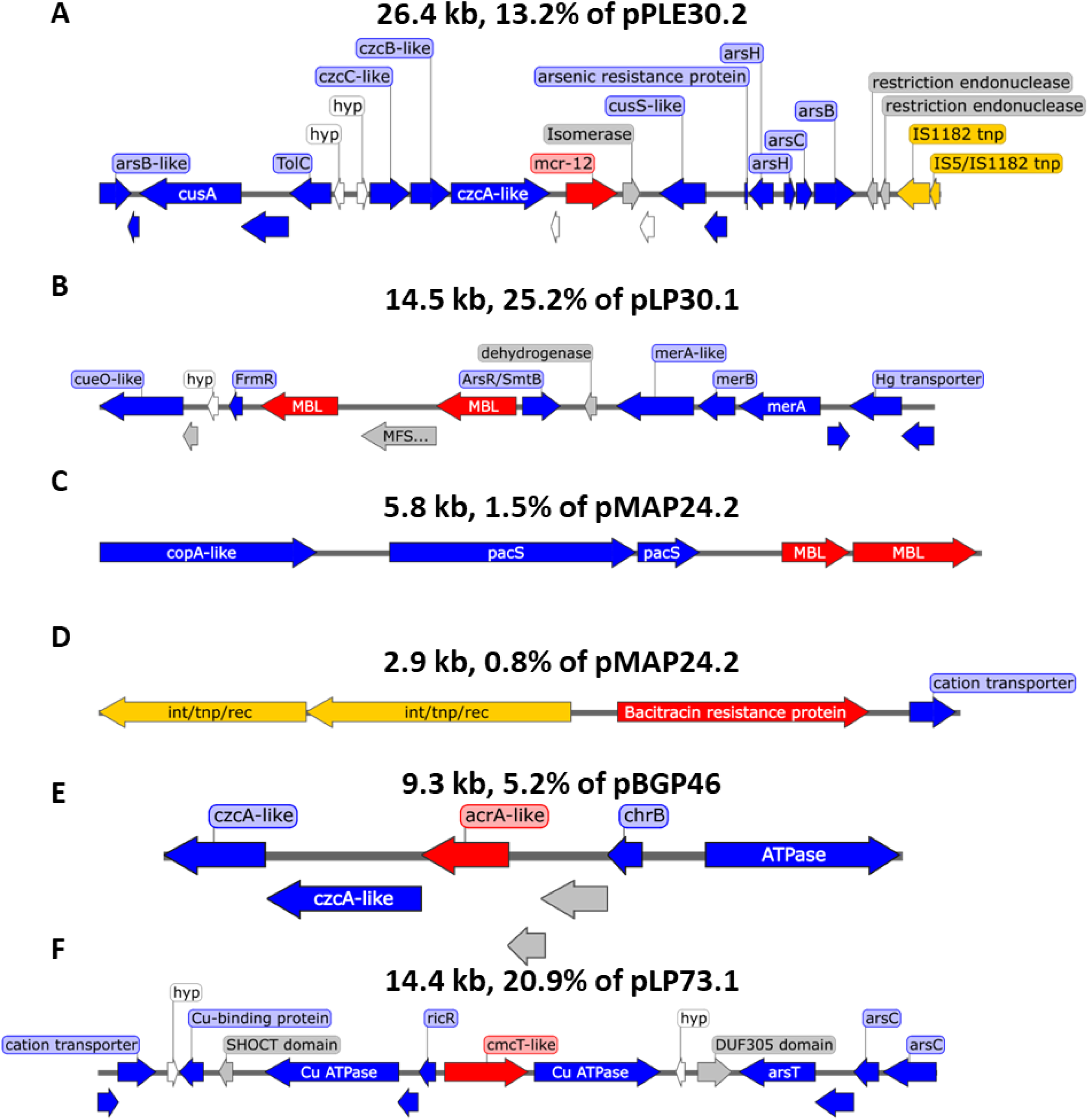
Resistance plasmid regions containing adjacent antibiotic resistance genes (ARG) and metal resistance genes (MRG) as evidence of co-resistance. Representative sequences from plasmids (**A**) pPLE30.2, (**B**) pLP30.1, (**C**, **D**) pMAP24.2, (**E**) pBGP46, (**F**) pLP73.1. Arrows indicate bioinformatically predicted genes, colour-coded by function: blue indicates MRGs, red indicates ARGs, orange represents mobile genetic elements, grey represents unrelated genes, and white genes of unknown identity.

*Leifsonia* sp. P30 also contained two resistance plasmids, with pLP30.1 providing the best evidence of co-resistance. The MOB_P_ type plasmid is 57.6 kb in size and carries a relaxase gene and a several T4SS structural components, though these do not appear to provide a complete system for conjugation. The plasmid possesses *cueO*-like and *ctpG*-like genes which encode copper oxidation and efflux, and *merRAB* which encodes mercury resistance. Two metallo-β-lactamase genes are situated between the *cueO*-like gene and the *mer* operon, indicative of co-resistance (Fig. 4B). Each resistance gene or operon is adjacent to separate regulator genes, making co-regulation unlikely.

*Mesorhizobium atlanticum* contained pMAP24.2, the second largest resistance plasmid in this study at 382 kb. The plasmid belongs to an undefined MOB type, containing a *virD2*-like relaxase gene, and potentially can be transferred by conjugation, as it carries a complete suite of *virB1–11* T4SS genes and a *virD4* T4CP gene. Copper resistance genes on pMAP24.2 include *cusAB*, *golT*, *mmcO*-like genes, *copA*-like genes, an *actP*-like gene, a *cueO*-like gene, and numerous miscellaneous copper oxidase, ATPase, and chaperone genes. Zinc resistance is encoded by multiple *zntA* genes and ZIP family transporter genes, while cadmium resistance is encoded by a miscellaneous cadmium ATPase gene. Nickel resistance is encoded by *nirA*-homologues, and mercury resistance is encoded by *merTPBAR*. Gene homologues of *czcD* encode resistance to cobalt, zinc, and cadmium. ARGs are located adjacent to MRGs, indicative of co-resistance. These include a metallo-β-lactamase gene adjacent to the *copA*-like gene and *pacS* (Fig. 4C), and a bacitracin resistance gene adjacent to a miscellaneous cation transporter gene (Fig. 4D). Two penicillin binding proteins, a *penA-*like gene and a *pbp3*-like gene, are present but not located near MRGs.

Another large plasmid, found in *B. guangxiense*, was named pBGP46 and is a 180.2 kb MOB_P_-type plasmid. This plasmid appears to be conjugative, since it contains the complete gene set for a VirB1–11 T4SS, VirD4 T4CP and VirD2 relaxase. MRGs include numerous *czcA* and *czc*-like cation efflux pump genes, the tellurium resistance gene *terB*, copper and silver-exporting ATPase genes, *cusAB*, and nickel and cobalt transporter genes, similar to *cnrA*. An *acrA*-like multidrug efflux pump gene is flanked downstream by a *czcA*-like gene, and upstream by *chrB*, which is consistent with co-resistance (Fig. 4E). The plasmid also contains a bacitracin resistance gene and a metallo-β-lactamase gene, though these are not near MRGs.

The final plasmid of interest for evidence of co-resistance, pLP73.1, is carried by *Leifsonia* sp. P73. This 68.7 kb MOB_P_-type plasmid harbours an *arsCT* operon, aquaglyceroporin gene, and *arsR* encoding arsenic resistance. The plasmid also carries numerous copper ATPase genes and a *czcD*-like gene. Notably, a *cmcT*-homologue, which encodes cephamycin export, is located directly between two copper ATPase genes, suggesting they are inherited as a single genetic entity (Fig. 4F). The *cmcT-*homologue is adjacent to a copper-sensing *ricR* regulator gene, which may regulate expression of these genes. The plasmid appears mobilisable, containing a *mobC* relaxase gene and a *virD4* coupling protein gene.

### The plasmid metagenome in industrial sediments reveals co-resistance, cross-resistance, and co-regulation plasmid features

The resistance plasmid analysis above is limited to plasmids found in culturable bacteria. To extend this characterisation to the entire bacterial community, the sediments were first treated with heavy metals for 28 days (2–20 mM cadmium or chromium) to enrich for cadmium or chromium resistant bacteria, and total DNA was then extracted directly from the bacterial fraction separated from the sediments. Plasmid DNA was selectively enriched by digestion with plasmid-safe DNAse, which specifically digests genomic DNA, and this was then amplified by multiple displacement amplification, sequenced, and assembled into 100–1000 contigs per sample (Table S7). These contigs were quite short (mean contig length ≤4 x mean read length), and only a minority of them were complete plasmids as indicated by circularisation. However, 37 contigs could be confidently assigned as plasmid sequences, since they carried plasmid-specific genes, including genes characteristic of six plasmid MOB and colicinogenic types (Table S8). Reference site sediment was excluded from this analysis, as no resistance plasmids had been detected in this sample in the isolate library.

Antibiotic or metal resistance genes were identified on 53 unique contigs, 20 of which were found at both industrial sites. Copper and silver MRGs were the most common resistance genes detected (Fig. 5), but arsenic, chromium, polymyxin, multi-antibiotic, and multi-metal resistance genes were also found. Of the 53 unique resistance contigs, 13 contained features of co-resistance (MRG and ARG on the same contig), cross-resistance (a gene that conveys both metal and antibiotic resistance), or co-regulation (one regulator adjacent to both metal and antibiotic genes) (Table S9). ARG and MRG combinations, which may indicate co-resistance, were detected on ten contigs. The *arnTFBCA* operon was detected on two contigs adjacent to a *cadA* cadmium resistance gene (Fig. S10A), which is notable as overexpression of the *arn* operon can confer polymyxin resistance [58]. A putative fluoroquinolone resistance operon, *emrAB*, was found near a putative *chrR* chromate reductase, along with an outer membrane porin that likely contributes to fluoroquinolone resistance (Fig. S10B). Similarly, an arsenic resistance operon *arsCBH* and an ArsR family regulator gene were co-localised with the multidrug efflux pump genes *acrAD* and *oprM* (Fig. S10C). Known cross-resistance genes were identified on seven contigs, including nine operons or isolated genes. These included homologues of the efflux pump *mdtABC* operon on four contigs (Fig. S10D), and two contigs carrying the gene for the inner membrane multidrug efflux pump CmeB (Fig. S10E), which exports fluoroquinolones, β-lactams, erythromycin, tetracycline, rifampicin, chloramphenicol, ampicillin, copper, and cobalt [59]. On another contig, a homologue of the cross-resistance gene *macAB* (conferring arsenite, macrolide, and penicillin resistance [13]) was co-located with *tolC* (Fig. S10F). Two contigs harboured multiple types of resistance genes resulting in both co-resistance and cross-resistance features. For example, an *mdtA* homologue, which encodes cross-resistance, was found adjacent to *acrB-tolC* efflux genes and a penicillin binding protein gene, *mrcA* (Fig. S10G), with class A and class D β-lactamase genes also present on the same contig. Additionally, a *bepG* homologue gene was found in proximity to *gesB* genes (Fig. S10H) which provides potential gold, phenicol, and β-lactam resistance [60]. This contig additionally carried a *silP* homologue 15.5 kb upstream.

**Fig. 5:**
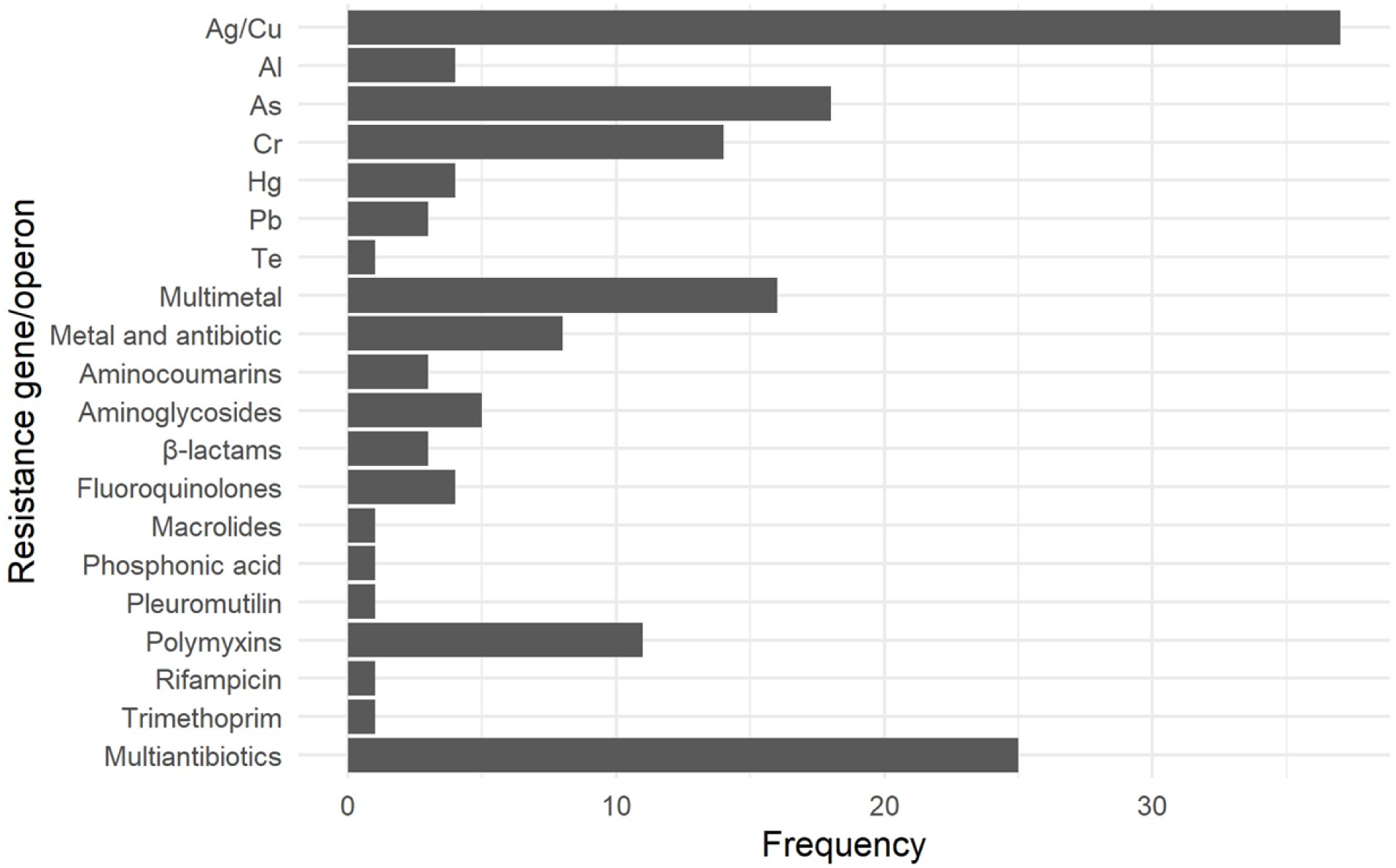
Copper, silver, arsenic, chromium, multi-metal, and multi-antibiotic resistance genes predominated in metagenomic plasmids. Metal and antibiotic resistance genes/operons detected across all assembled contigs. Multiple genes predicted to belong to the same hypothetical operon are represented as a single operon unit.

## Discussion

Co-selection is a major driver of AMR, and the link between heavy metal exposure and selection for ARGs has been well-established [3, 7, 8]. The present study explores the broader question of how heavy metals shape co-selection for antibiotic resistance in natural environments. The broad range of culturable metal-resistant bacteria were found to harbour a diverse and novel set of plasmids that featured predicted MRGs and ARGs. These resistance plasmids were exclusively found in isolates cultured from sediments impacted by coal ash or smelting and were not present in sediment from an uncontaminated reference site nearby, supporting the hypothesis that these resistance plasmids play a role in the bacterial response to industrial heavy metal stress. Two-thirds of the metal-resistance plasmids that were found in these isolates harboured both ARGs and MRGs on the plasmid, providing strong evidence for co-resistance. A complementary, culture-independent analysis of plasmids present in the sediment metagenomes revealed an even greater diversity of plasmid types, with many examples of co-located MRGs and ARGs and several examples of plasmid-borne efflux pump genes that may contribute to cross-resistance. Unexpectedly, there was only limited evidence for the transfer of these plasmid-borne resistance genes by transposons. The environmental hosts for most of the plasmids studied do not pose a risk to humans, although *Pantoea* may act as an opportunistic pathogen [61]. It is therefore of considerable interest to determine whether the plasmids in this collection can disseminate resistance genes into other bacteria, and ultimately into relevant human pathogens.

### Impact of the sediment’s physicochemical characteristics on the bacterial community

The sampling locations used in this study were selected to overlap with heavily contaminated sites that have been investigated previously. Although the concentrations of heavy metals were found to be elevated in comparison to the reference site, they were generally several fold lower than in previous reports [24, 25, 62]. This appears to suggest a reduction in metal load at the site over time, but significant heterogeneity in heavy metal concentrations has been previously noted at Site P [62], and it is therefore difficult to determine whether the observed differences are due to spatial or temporal factors.

Despite the heterogeneity of the sediment, the findings reported here are consistent with the hypothesis that a higher heavy metal load drives bacterial selection for MRGs and for the MGEs that carry them. This was supported by the greater abundance of cadmium-and copper-resistant cultivable bacteria (Fig. 1) at the industrial sites compared to the control site. This pattern did not extend to zinc resistance, but this may reflect the relatively high zinc levels at Site L (Fig. S1), and the fact that zinc is generally more abundant in natural environments than copper or cadmium [63]. Contamination of the sediments with mercury, arsenic, and chromium likely had a greater biological impact than copper or cadmium, as indicated by the NIRI risk index, and served as a selection pressure on the microbial community. Mercury levels in particular were significantly correlated with the number of cultivable bacteria resistant to copper and cadmium (Fig. S2). Arsenic [13, 64], chromium [65, 66], and mercury [11, 67] have all been associated with co-selection for ARGs, and further work is needed to explore the effect of metal load on the development of resistance to these elements.

Sites impacted by coal ash or smelting generally low in microbial diversity [23, 68], and the current study also found that the sediment of Site E, which is impacted by coal ash, had lower diversity than the control sediment (Fig. S4). Site E had similar diversity in culturable metal-resistant isolates to the control site consistent with the enrichment for metal-resistant taxa. Metal-resistant bacteria previously isolated from coal ash include mercury-resistant and chromium-resistant *Micrococcus, Staphylococcus*, and *Bacillus* [69, 70], but of these, only metal-resistant *Bacillus* was isolated in the current study. The sediment from the lead-zinc smelter at Site P was dominated by *Actinomycetota*, which contrasts with previous assessments of soil impacted by lead-zinc smelter emissions, which contained a high abundance of *Pseudomonadota* [68, 71].

Resistance plasmids were detected in only 13 of the 64 sequenced metal-resistant isolates, which suggests that the majority of resistance genes in these isolates were chromosomally derived. However, the absence of resistance plasmids from the control site is noteworthy and supports the theory that metal contamination exerted a significant selection pressure in the industrial sediments. A limitation of this study, however, is that linear plasmids, which are common in *Actinomycetota*, would not have been detected, as they are not circularised [72].

Given the contamination of the sediment by heavy metals, it is unsurprising that MRGs were an order of magnitude more abundant than ARGs among the plasmids from isolates and in the plasmid metagenome. Copper, zinc, and cadmium resistance genes dominated the MRG types, as expected given the selection for copper-, zinc-, or cadmium-resistant bacteria, and cadmium-resistant communities. However, arsenic resistance was also relatively common in plasmids from both culturable and metagenomic communities (Fig. 3, 5), which may reflect arsenic being a notable pollutant at both industrial sites (Table 1). Mercury resistance genes were not as common as in previous plasmid metagenome studies focused on groundwater samples near a uranium processing facility [73] or metal-enriched high-altitude lakes [74]. At these sites, less than 5% of the total ORFs were assigned as MRGs or ARGs [73, 74], which is lower than the proportions in the plasmids annotated in the current study, but similar MRGs predominated (zinc, copper, arsenic, multi-metal resistance genes).

The presence of ARGs in these plasmids is notable, given the assumed lack of antibiotic selection. Antibiotic residues were not measured in the collected samples, as there were no known agricultural, industrial, municipal, or clinical sources upstream of the sites. The most common group of ARGs in the plasmid metagenome were multi-antibiotic efflux pump genes, which are agents of cross-resistance [13, 60]. Many of these multi-antibiotic resistance genes fall into the highest ARG risk quartile defined by Zhang et al [75]. Although functionality for these genes were not assessed here, it is known that some of these genes are not always involved in antibiotic resistance, since *acrB*, *acrD* and *ceoB* are also involved in phosphorus cycling, and *macB* plays a role in sulfur cycling [75]. Some resistance operons appeared to be incomplete on the plasmid sequence, such as the *mdtABC* or *cmeB* examples. These may be ‘orphan genes’, or the missing components (*tolC* and *cmeA*/*cmeC* in these examples) may be located on the chromosome or on the plasmid beyond the point of contig truncation. The regulator genes adjacent to the putative resistance genes are predicted to be involved in the resistance gene expression, but this remains to be confirmed.

### New plasmid-borne metal and antibiotic resistance profiles

Resistance to multiple metals were confirmed to be plasmid-linked in *P. litoralis* E30 and *S. hankookensis* E10. The increased susceptibility of plasmid-cured isolates strongly supports that zinc, cobalt, and cadmium resistance is mediated by *czcCBA* and *czcD*-homologues on pPLE30.2, and *czcCBAP* and *dmeF* on pSHE10.1 (Table 3). Plasmid pSHE10.1 is the likelier carrier of functional cobalt-zinc-cadmium resistance genes as it harbours the complete *czcCBA* efflux pump operon and *czcP*, whereas pSHE10.2, contains only *czcA*. Similarly, arsenite and arsenate resistance were attributed to *arsHCB* on pPLE30.2 and *arsCBNH* on pSHE10.1; mercury resistance with *merTPA* on pSHE10.1; and nickel resistance to either *dmeF* on pSHE10.1 or *nirAB*- and *nccBC-*homologues on pSHE10.2. Copper resistance was also linked to pSHE10.1 which carries both *cop* and *cus* operons. In contrast, the loss of *cusAB* and *copABCDG-*homologue operons from pPLE30.2 in plasmid-cured *P. litoralis* did not alter its copper susceptibility compared to the wild-type. This does not exclude a role for these genes in copper resistance, as their absence may be compensated by chromosomal homeostasis genes essential for maintaining redox-active enzymes [76].

Unfortunately, confirming the resistance phenotype by plasmid curing is often difficult, since the efficiency of the acridine orange method varies significantly between different bacterial species and plasmid types, especially for large plasmids [77]. Many of the plasmids studied here also carried toxin/antitoxin systems, which may have hindered curing attempts. Rather than removing the plasmid entirely, targeted knock-out, overexpression, or heterologous expression of the resistance genes can be used to further examine the function of these genes [78, 79].

In addition to the plasmid-borne heavy metal resistance phenotypes, *S. hankookensis* exhibited streptomycin resistance, consistent with Yoon et al [80]. Aminoglycoside and β-lactam resistance in *Sphingomonas* is intrinsic [81], suggesting that the antibiotic target is resistant or that the relevant ARGs are carried on the chromosome. The current study offers the first characterisation of heavy metal and antibiotic resistance phenotypes for *M. atlanticum* and *Amnibacterium kyonggiense*, and builds on the first resistance characterisation of *P. litoralis* [57] with the inclusion of neomycin resistance (Table S5). *P. litoralis* E30 was polymyxin resistant due to carriage of the novel *mcr-12* gene on plasmid pPL30.2, demonstrated by the substantial reduction in polymyxin B MIC for the plasmid-cured *P. litoralis* strain (Table 3).

Many of the plasmids identified in sediment isolates also contained genes annotated as metallo-β-lactamase genes, since they shared the αββα-fold domain of the metallo-β-lactamase superfamily [82]. It must be noted, however, that many members of this superfamily have functions that are unrelated to antibiotic resistance [82], and indeed none of the predicted β-lactamase genes in the current study appear to encode a functional β-lactamase under the *in vitro* conditions used.

The inability to verify functional β-lactamases reinforces the fact that homology-based methods of classifying genes must be applied with care, since many of the ‘resistance’ gene products in CARD and AMRFinder are in fact alleles of housekeeping genes, which confer resistance as a consequence of a specific point mutation (e.g. the DNA gyrase gene *gyrA,* where a single nucleotide polymorphism can result in quinolone resistance [83]).

Alternatively, they may be transcriptional regulators of functional resistance genes, but do not themselves encode resistance functions.

### Co-resistance and horizontal gene transfer

Co-resistance is generally thought to be more common than cross-resistance in mediating co-selection [8], and the most frequent form of co-resistance is the presence of both MRGs and ARGs on the same plasmid or transposon [7]. The resistance plasmids identified in the present study may enable the spread of resistance genes across phylogenetically divergent bacterial groups. Thirteen complete plasmids were found to carry both ARGs and MRGs, and while there were some indications of genes for cross-resistance mechanisms and genetic arrangements consistent with co-regulation, these were less frequent than the clear observed patterns of co-resistance. Five plasmids and ten metagenomic plasmid contigs had different combinations of ARGs located close to MRGs (less than 1 kb apart), including *mcr-12*, *arnTFBCA*, *rosB*, *macAB*, *bacA*, *cmcT*, β-lactamase resistance genes, *acrA*, *czcCBA*, *cadA*, *chrB*, and copper resistance genes (Fig. 4, S10). The close proximity of these gene combinations makes them unlikely to be separated during recombination events, thus selection for one resistance gene would indirectly lead to the acquisition of the other. This is especially clear for the plasmids identified within isolates, but for the plasmid metagenome sequences some care in interpretation is required, since the phi29-mediated multiple displacement amplification used here can generate chimeric sequences from regions of homology only 6–12 bp in length [84]. This does not undermine the detection of resistance genes but may result in the duplication of such genes in the assembly. Flye v2.6 has previously been evaluated as the most reliable and robust Oxford Nanopore sequence assembly tool for plasmids, especially at the low read depth relevant to the metagenomic data of this study, but it tends to struggle with plasmid circularisation [85]. The version used here (v2.9.3) is an improved version but appears still prone to this error.

The typed plasmids belonged primarily to MOB_P_ or MOB_F_ types (Fig. S6). Both types are common carriers of ARGs, with extended spectrum β-lactamases often carried by MOB_F_ plasmids [15]. MOB_F_ plasmids are confined to *Enterobacteriaceae*, however some possess multiple replicon sequences that allow replication in more diverse taxa [86]. MOB_P_ plasmids are found in diverse taxa belonging to *Pseudomonadota* and *Actinomycetota* [87]. MOB_P_ plasmids are often conjugative [15] and several of the MOB_P_ plasmids reported here contained the relaxase, T4CP, and T4SS genes required for conjugation (Table 2). No *oriT* sequences were detected by the oriTfinder algorithm [42], but it is likely that these plasmids have a diverse *oriT* not predicted by the current algorithm, which is trained on conserved inverted repeat sequences flanking the *nic* site in *Gammaproteobacteria* plasmids. The potential for mobility needs to be further confirmed using biparental and triparental matings to distinguish between conjugative and mobilisable plasmids, but for several resistance plasmids from different isolates it appears that this has already occurred, since they shared large regions of homology.

Intra-cell gene mobilisation events are often indicated by the presence of transposons. Almost every plasmid in this study carried transposase genes, with the most common being Tn*3*-type transposases (Table S6), which are often associated with ARGs [15]. No complete transposon was found to carry a resistance gene, but in several cases transposase genes were directly adjacent to β-lactamase or copper oxidase genes (Fig. S9). This has been observed before in environmental resistance plasmids [7], and it has been suggested that the presence of transposase genes within 5 kb of resistance genes is an indicator of past transposition events [75]. Many of the transposons in this study were found to lack direct or inverted flanking repeat sequences, and they may therefore have lost the ability to transpose [15], however, it remains possible that some resistance phenotypes in this study may be expressed from a promoter in an adjacent transposon or IS. This has been seen before in *Acinetobacter baumannii,* where cephalosporin resistance results from the insertion of IS*Aba1* upstream of the endogenous *ampC* gene [88].

Co-selection may also be due to cross-resistance or co-regulation [8]. The frequency of *mdtABC* homologues in the plasmid metagenome, encoding multi-substrate efflux pumps, suggests that cross-resistance mechanisms are present in these sediments (Fig. S10). However, the encoded proteins share only 30–70% amino acid identity with known resistance proteins, and therefore they may represent novel efflux pumps with different substrate ranges. Little can be inferred from our genomic data about the presence of co-regulation, particularly for co-regulated genes that are not also adjacent. There were four instances where plasmid-borne ARGs were adjacent to MRGs with only one regulator gene within the immediate 5 kb flanking sequence, but it requires considerable further work to establish whether these genes are part of the same regulon. The involvement of regulators encoded *in trans* is also possible but determining such relationships based purely on DNA sequence is challenging, and requires co-expression studies.

## Conclusion

This study highlights freshwater sediments impacted by sub-MIC concentrations of metals as hotspots for the co-selection of MRGs and ARGs. Not only do these sites pose a biological risk due to the innate toxicity of heavy metals, but the selection pressure of heavy metals could, in part, explain the persistence of AMR despite improved antimicrobial stewardship. It is recommended that a management approach focused on minimisation be applied to the use of heavy metals. The environmental ARG reservoir is far larger than the clinical reservoir, and the environmental origin of many clinical resistance genes demonstrates that these reservoirs intersect likely facilitated by HGT. The presence of large multi-resistance potentially mobile plasmids places a profound importance on cataloguing the environmental resistome and evaluating the risk to public health in locations such as metal-impacted freshwater sediment.

## Acknowledgements

We thank Mehrad Hamidian (University of Technology Sydney) and Michael Gillings (Macquarie University) for their recommendations on the project scope. We extend our thanks to Mark Somerville and Scott Mitchell (The University of Sydney) for their technical support in bacterial strain collation and plasmid characterisation.

## Authors contributions

BFG, NVC, AKC, RNZ, and MAK conceptualised and planned the project. BFG performed the experiments, apart from ion chromatography which was conducted by MT who also provided technical assistance in bioinformatic pipelines. BFG wrote the original manuscript and produced the tables and figures with assistance from MAK. All authors reviewed and approved the final manuscript.

## Data availability statement

Assembled genomes for isolates and annotated sequences for all identified plasmids are available in GenBank, and accession numbers are provided in Table S2 (isolates) and Table 2 (plasmids). Raw pooled sequence reads are available as SRA accession PRJNA1263914.

## Ethical approval

This article does not contain any studies with human participants or animals performed by any of the authors.

## References

1. Naghavi M, Vollset SE, Ikuta KS et al. Global burden of bacterial antimicrobial resistance 1990-2021: a systematic analysis with forecasts to 2050. Lancet. 2024;404:1199–226 10.1016/S0140-6736(24)01867-1

2. Larsson DGJ, Flach CF. Antibiotic resistance in the environment. Nat Rev Microbiol. 2022;20:257–69 10.1038/s41579-021-00649-x

3. Vats P, Kaur UJ, Rishi P. Heavy metal-induced selection and proliferation of antibiotic resistance: a review. J Appl Microbiol. 2022;132:4058–76 10.1111/jam.15492

4. Arana DM, Saez D, García-Hierro P et al. Concurrent interspecies and clonal dissemination of OXA-48 carbapenemase. Clin Microbiol Infect. 2015;21:e1–4 10.1016/j.cmi.2014.07.008

5. Feng YJ. Transferability of MCR-1/2 polymyxin resistance: complex dissemination and genetic mechanism. ACS Infect Dis. 2018;4:291–300 10.1021/acsinfecdis.7b00201

6. Bernier SP, Surette MG. Concentration-dependent activity of antibiotics in natural environments. Front Microbiol. 2013;4:20 10.3389/fmicb.2013.00020

7. Baker-Austin C, Wright MS, Stepanauskas R et al. Co-selection of antibiotic and metal resistance. Trends Microbiol. 2006;14:176–82 10.1016/j.tim.2006.02.006

8. Gillieatt BF, Coleman NV. Unravelling the mechanisms of antibiotic and heavy metal resistance co-selection in environmental bacteria. FEMS Microbiol Rev. 2024;48:fuae017 10.1093/femsre/fuae017

9. Flach CF, Pal C, Svensson CJ et al. Does antifouling paint select for antibiotic resistance? Sci Total Environ. 2017;590-591:461–68 10.1016/j.scitotenv.2017.01.213

10. Vignaroli C, Pasquaroli S, Citterio B et al. Antibiotic and heavy metal resistance in enterococci from coastal marine sediment. Environ Pollut. 2018;237:406–13 10.1016/j.envpol.2018.02.073

11. Pal C, Bengtsson-Palme J, Kristiansson E et al. Co-occurrence of resistance genes to antibiotics, biocides and metals reveals novel insights into their co-selection potential. BMC Genom. 2015;16:964 10.1186/s12864-015-2153-5

12. Li LG, Xia Y, Zhang T. Co-occurrence of antibiotic and metal resistance genes revealed in complete genome collection. ISME J. 2017;11:651–62 10.1038/ismej.2016.155

13. Shi K, Cao M, Li C et al. Efflux proteins MacAB confer resistance to arsenite and penicillin/macrolide-type antibiotics in *Agrobacterium tumefaciens* 5A. World J Microbiol Biotechnol. 2019;35:115 10.1007/s11274-019-2689-7

14. Slifierz MJ, Friendship RM, Weese JS. Methicillin-resistant *Staphylococcus aureus* in commercial swine herds is associated with disinfectant and zinc usage. Appl Environ Microbiol. 2015;81:2690–5 10.1128/aem.00036-15

15. Partridge SR, Kwong SM, Firth N et al. Mobile genetic elements associated with antimicrobial resistance. Clin Microbiol Rev. 2018;31:e00088–17 10.1128/cmr.00088-17

16. Virolle C, Goldlust K, Djermoun S et al. Plasmid transfer by conjugation in Gram-negative bacteria: from the cellular to the community level. Genes. 2020;11:1239 10.3390/genes11111239

17. Million-Weaver S, Camps M. Mechanisms of plasmid segregation: have multicopy plasmids been overlooked? Plasmid. 2014;75:27–36 10.1016/j.plasmid.2014.07.002

18. McArthur AG, Waglechner N, Nizam F et al. The comprehensive antibiotic resistance database. Antimicrob Agents Chemother. 2013;57:3348–57 10.1128/aac.00419-13

19. Feldgarden M, Brover V, Gonzalez-Escalona N et al. AMRFinderPlus and the Reference Gene Catalog facilitate examination of the genomic links among antimicrobial resistance, stress response, and virulence. Sci Rep. 2021;11:12728 10.1038/s41598-021-91456-0

20. Pal C, Bengtsson-Palme J, Rensing C et al. BacMet: antibacterial biocide and metal resistance genes database. Nucleic Acids Res. 2014;42:D737–D43 10.1093/nar/gkt1252

21. Bondarczuk K, Markowicz A, Piotrowska-Seget Z. The urgent need for risk assessment on the antibiotic resistance spread via sewage sludge land application. Environ Int. 2016;87:49–55 10.1016/j.envint.2015.11.011

22. Stepanauskas R, Glenn TC, Jagoe CH et al. Elevated microbial tolerance to metals and antibiotics in metal-contaminated industrial environments. Environ Sci Technol. 2005;39:3671–78 10.1021/es048468f

23. Thomas JC, Oladeinde A, Kieran TJ et al. Co-occurrence of antibiotic, biocide, and heavy metal resistance genes in bacteria from metal and radionuclide contaminated soils at the Savannah River Site. Microb Biotechnol. 2020;13:1179–200 10.1111/1751-7915.13578

24. Winn P, Lynch J, Woods G. NSW water pollution and our aging coal-fire power stations. Report. Centre HCE, 2020 https://www.hcec.org.au/out-of-the-ashes-ii

25. Harvey PJ, Mabbott R, Rouillon M et al. Delineating the spatial extent of smelter-related atmospheric fallout using a rapid assessment technique. Appl Geochem. 2018;96:35–41 10.1016/j.apgeochem.2018.06.003

26. Varhammar A, McLean CM, Yu RMK et al. Uptake and partitioning of metals in the Australian saltmarsh halophyte, samphire (*Sarcocornia quinqueflora*). Aquat Bot. 2019;156:25–37 10.1016/j.aquabot.2019.04.001

27. Li Y, Chen H, Song L et al. Effects on microbiomes and resistomes and the source-specific ecological risks of heavy metals in the sediments of an urban river. J Hazard Mater. 2020;409:124472 10.1016/j.jhazmat.2020.124472

28. Chen J, Li J, Zhang H et al. Bacterial heavy-metal and antibiotic resistance genes in a copper tailing dam area in Northern China. Front Microbiol. 2019;10:1916 10.3389/fmicb.2019.01916

29. Hakanson L. An ecological risk index for aquatic pollution control. A sedimentological approach Water Res. 1980;14:975–1001 10.1016/0043-1354(80)90143-8

30. Callahan BJ, McMurdie PJ, Rosen MJ et al. DADA2: high-resolution sample inference from Illumina amplicon data. Nat Methods. 2016;13:581–83 10.1038/nmeth.3869

31. McMurdie PJ, Holmes S. phyloseq: an R package for reproducible interactive analysis and graphics of microbiome census data. PLoS One. 2013;8:e61217 10.1371/journal.pone.0061217

32. Quast C, Pruesse E, Yilmaz P et al. The SILVA ribosomal RNA gene database project: improved data processing and web-based tools. Nucleic Acids Res. 2012;41:D590–96 10.1093/nar/gks1219

33. Coleman NV, Holmes AJ. The native *Pseudomonas stutzeri* strain Q chromosomal integron can capture and express cassette-associated genes. Microbiol-SGM. 2005;151:1853–64 10.1099/mic.0.27854-0

34. Koeuth T, Versalovic J, Lupski JR. Differential subsequence conservation of interspersed repetitive *Streptococcus pneumoniae* BOX elements in diverse bacteria. Genome Res. 1995;5:408–18 10.1101/gr.5.4.408

35. Tamura K, Stecher G, Kumar S. MEGA11: Molecular Evolutionary Genetics Analysis version 11. Mol Biol Evol. 2021;38:3022–27 10.1093/molbev/msab120

36. Wick R Filtlong [Computer software]. Github. https://github.com/rrwick/Filtlong. 2017.

37. De Coster W Nanoplot [Computer software]. Github. https://github.com/wdecoster/NanoPlot. 2018.

38. Kolmogorov M, Yuan J, Lin Y et al. Assembly of long, error-prone reads using repeat graphs. Nature Biotechnol. 2019;37:540–46 10.1038/s41587-019-0072-8

39. Seemann T Barrnap 0.9: rapid ribosomal RNA prediction [Computer software]. Github. https://github.com/tseemann/barrnap. 2020.

40. Carattoli A, Zankari E, García-Fernández A et al. *In silico* detection and typing of plasmids using PlasmidFinder and plasmid multilocus sequence typing. Antimicrob Agents Chemother. 2014;58:3895–903 10.1128/aac.02412-14

41. Garcillán-Barcia MP, Redondo-Salvo S, Vielva L et al. MOBscan: automated annotation of MOB relaxases. Methods Mol Biol. 2020;2075:295–308 10.1007/978-1-4939-9877-7_21

42. Li X, Xie Y, Liu M et al. OriTfinder: a web-based tool for the identification of origin of transfers in DNA sequences of bacterial mobile genetic elements. Nucleic Acids Res. 2018;46:W229–34 10.1093/nar/gky352

43. Seemann T. Prokka: rapid prokaryotic genome annotation. Bioinformatics. 2014;30:2068–69 10.1093/bioinformatics/btu153

44. Siguier P, Perochon J, Lestrade L et al. ISfinder: the reference centre for bacterial insertion sequences. Nucleic Acids Res. 2006;34:D32–D36 10.1093/nar/gkj014

45. Warburton PE, Giordano J, Cheung F et al. Inverted repeat structure of the human genome: The X-chromosome contains a preponderance of large, highly homologous inverted repeats that contain testes genes. Genome Res. 2004;14:1861–9 10.1101/gr.2542904

46. Wishart DS, Han S, Saha S et al. PHASTEST: faster than PHASTER, better than PHAST. Nucleic Acids Res. 2023;51:W443–W50 10.1093/nar/gkad382

47. 47. Bell SM, Pham JN, Rafferty DL et al. Antibiotic susceptibility testing by the CDS method: A manual for medical and veterinary laboratories Report. Services SEAL, 2016 https://cdstest.net/wordpress/wp-content/uploads/Antibiotic-Susceptibility-Testing-by-the-CDS-Method-8th-Edition.pdf

48. The European Committee on Antimicrobial Susceptibility Testing. Breakpoint tables for interpretation of MICs and zone diameters. http://www.eucast.org (03 February 2025, date last accessed).

49. Livermore DM, Brown DFJ. Detection of β-lactamase-mediated resistance. J Antimicrob Chemother. 2001;48:59–64 10.1093/jac/48.suppl_1.59

50. Heringa S, Monroe J, Herrick J. A simple, rapid method for extracting large plasmid DNA from bacteria. Nat Preced. 2007 10.1038/npre.2007.1249.1

51. Ricker N, Spoja BS, May N et al. Incorporating the plasmidome into antibiotic resistance surveillance in animal agriculture. Plasmid. 2021;113:102529 10.1016/j.plasmid.2020.102529

52. Dixon P. VEGAN, a package of R functions for community ecology. J Veg Sci. 2003;14:927–30 10.1111/j.1654-1103.2003.tb02228.x

53. Wickham H. ggplot2. Elegant graphics for data analysis, 2 edn.: Springer Cham, 2016.

54. Nies DH. The cobalt, zinc, and cadmium efflux system CzcABC from *Alcaligenes eutrophus* functions as a cation-proton antiporter in *Escherichia coli*. J Bacteriol. 1995;177:2707–12 10.1128/jb.177.10.2707-2712.1995

55. Nishino K, Nikaido E, Yamaguchi A. Regulation of multidrug efflux systems involved in multidrug and metal resistance of *Salmonella enterica* serovar typhimurium. J Bacteroiol. 2007;189:9066–75 10.1128/jb.01045-07

56. Wang D, Fierke CA. The BaeSR regulon is involved in defense against zinc toxicity in *E. coli*. Metallomics. 2013;5:372–83 10.1039/c3mt20217h

57. Gillieatt BF, Maharjan RP, Cain JA et al. Novel polymyxin resistance gene family mcr-12 from environmental Pigmentiphaga litoralis. Nat Commun. 2025;**under revision**

58. Leung LM, Cooper VS, Rasko DA et al. Structural modification of LPS in colistin-resistant, KPC-producing *Klebsiella pneumoniae*. J Antimicrob Chemother. 2017;72:3035–42 10.1093/jac/dkx234

59. Lin J, Michel Linda O, Zhang Q. CmeABC functions as a multidrug efflux system in *Campylobacter jejuni*. Antimicrob Agents Chemother. 2002;46:2124–31 10.1128/aac.46.7.2124-2131.2002

60. Conroy O, Kim EH, McEvoy MM et al. Differing ability to transport nonmetal substrates by two RND-type metal exporters. FEMS Microbiol Lett. 2010;308:115–22 10.1111/j.1574-6968.2010.02006.x

61. Dutkiewicz J, Mackiewicz B, Lemieszek MK et al. *Pantoea agglomerans*: a mysterious bacterium of evil and good. Part III. Deleterious effects: infections of humans, animals and plants. 2016;23:197–205 10.5604/12321966.1203878

62. Kim KR, Owens G, Naidu R. Heavy metal distribution, bioaccessibility, and phytoavailability in long-term contaminated soils from Lake Macquarie, Australia. Aust J Soil Res. 2009;47:166–76 10.1071/sr08054

63. Hu Z, Gao S. Upper crustal abundances of trace elements: a revision and update. Chem Geol. 2008;253:205–21 10.1016/j.chemgeo.2008.05.010

64. Zhao X, Shen JP, Zhang LM et al. Arsenic and cadmium as predominant factors shaping the distribution patterns of antibiotic resistance genes in polluted paddy soils. J Hazard Mater. 2020;389:121838 10.1016/j.jhazmat.2019.121838

65. Hubeny J, Harnisz M, Korzeniewska E et al. Industrialization as a source of heavy metals and antibiotics which can enhance the antibiotic resistance in wastewater, sewage sludge and river water. PLoS One. 2021;16:e0252691 10.1371/journal.pone.0252691

66. Gupta S, Graham DW, Sreekrishnan TR et al. Heavy metal and antibiotic resistance in four Indian and UK rivers with different levels and types of water pollution. Sci Total Environ. 2023;857:159059 10.1016/j.scitotenv.2022.159059

67. Wireman J, Liebert CA, Smith T et al. Association of mercury resistance with antibiotic resistance in the Gram-negative fecal bacteria of primates. Appl Environ Microbiol. 1997;63:4494–503 10.1128/aem.63.11.4494-4503.1997

68. Li S, Zhao B, Jin M et al. A comprehensive survey on the horizontal and vertical distribution of heavy metals and microorganisms in soils of a Pb/Zn smelter. J Hazard Mater. 2020;400:123255 10.1016/j.jhazmat.2020.123255

69. Ghosh S, Mukherjee P, Pati P et al. Isolation and characterization of mercury resistant bacteria from fly ash sample of Mejia Thermal Power Plant, W. B, India for application in bioremediation and phytoremediation. IOSR J Environ Sci Toxicol Food Technol. 2015;9:70–76 10.9790/2402-09717076

70. Roychowdhury R, Mukherjee P, Roy M. Identification of chromium resistant bacteria from dry fly ash sample of Mejia MTPS Thermal Power Plant, West Bengal, India. Bull Environ Contam Toxicol. 2016;96:210–16 10.1007/s00128-015-1692-4

71. Yang F, Zhang FL, Li HP et al. Contribution of environmental factors on the distribution of antibiotic resistance genes in agricultural soil. Eur J Soil Biol. 2021;102:103269 10.1016/j.ejsobi.2020.103269

72. Dib JR, Wagenknecht M, Farias ME et al. Strategies and approaches in plasmidome studies uncovering plasmid diversity disregarding of linear elements? Front Microbiol. 2015;6:463 10.3389/fmicb.2015.00463

73. Kothari A, Wu YW, Chandonia JM et al. Large circular plasmids from groundwater plasmidomes span multiple incompatibility groups and are enriched in multimetal resistance genes. mBio. 2019;10:e02899–18 10.1128/mBio.02899-18

74. Perez MF, Kurth D, Farias ME et al. First report on the plasmidome from a high-altitude lake of the Andean Puna. Front Microbiol. 2020;11:1343 10.3389/fmicb.2020.01343

75. Zhang Z, Zhang Q, Wang T et al. Assessment of global health risk of antibiotic resistance genes. Nat Commun. 2022;13:1553 10.1038/s41467-022-29283-8

76. Nies DH. Microbial heavy-metal resistance. Appl Microbiol Biotechnol. 1999;51:730-50 10.1007/s002530051457

77. Trevors JT. Plasmid curing in bacteria. FEMS Microbiol Rev. 1986;1:149–57 10.1111/j.1574-6968.1986.tb01189.x

78. Hayashi S, Abe M, Kimoto M et al. The DsbA-DsbB disulfide bond formation system of *Burkholderia cepacia* is involved in the production of protease and alkaline phosphatase, motility, metal resistance, and multi-drug resistance. Microbiol Immunol. 2000;44:41–50 10.1111/j.1348-0421.2000.tb01244.x

79. Deveson Lucas D, Crane B, Wright A et al. Emergence of high-level colistin resistance in an *Acinetobacter baumannii* clinical isolate mediated by inactivation of the global regulator H-NS. Antimicrob Agents Chemother. 2018;62:e02442–17 10.1128/aac.02442-17

80. Yoon J-H, Park S, Kang S-J et al. *Sphingomonas hankookensis* sp. nov., isolated from wastewater. Int J Syst Evol Microbiol. 2009;59:2788–93 10.1099/ijs.0.008680-0

81. Vaz-Moreira I, Nunes OC, Manaia CM. Bacterial diversity and antibiotic resistance in water habitats: searching the links with the human microbiome. FEMS Microbiol Rev. 2014;38:761–78 10.1111/1574-6976.12062

82. Daiyasu H, Osaka K, Ishino Y et al. Expansion of the zinc metallo-hydrolase family of the β-lactamase fold. FEBS Lett. 2001;503:1–6 10.1016/S0014-5793(01)02686-2

83. Bengtsson-Palme J, Larsson DGJ, Kristiansson E. Using metagenomics to investigate human and environmental resistomes. J Antimicrob Chemother. 2017;72:2690–703 10.1093/jac/dkx199

84. Nelson JR. Random-primed, Phi29 DNA polymerase-based whole genome amplification. Curr Protoc Mol Biol. 2014;105:Unit 15.13. 10.1002/0471142727.mb1513s105

85. Wick RR, Holt KE. Benchmarking of long-read assemblers for prokaryote whole genome sequencing. F1000Res. 2019;8:2138 10.12688/f1000research.21782.1

86. Villa L, García-Fernández A, Fortini D et al. Replicon sequence typing of IncF plasmids carrying virulence and resistance determinants. J Antimicrob Chemother. 2010;65:2518–29 10.1093/jac/dkq347

87. Redondo-Salvo S, Fernández-López R, Ruiz R et al. Pathways for horizontal gene transfer in bacteria revealed by a global map of their plasmids. Nat Commun. 2020;11:3602 10.1038/s41467-020-17278-2

88. Héritier C, Poirel L, Nordmann P. Cephalosporinase over-expression resulting from insertion of IS*Aba1* in *Acinetobacter baumannii*. Clin Microbiol Infect. 2006;12:123–30 10.1111/j.1469-0691.2005.01320.x

