## Supplementary Information for "The role of heavy metals in the co-selection of plasmid-borne metal and antibiotic resistance genes from industrially contaminated sediments"

**Table S1: Primers.** Primers labelled "BFG" were specifically designed for this study.

| Primer | Sequence (5' – 3') | Purpose |
|---|---|---|
| Random Hexamer | NNNN⁺N⁺N | Multiple displacement amplification |
| BOX-A1R | CTACGGCAAGGCGACGCTGACG | BOX PCR |
| 341F | CCTACGGGNGGCWGCAG | 16S V3 |
| 805R | GACTACHVGGGTATCTAATCC | 16S V4 |
| 806R | GGACTACNVGGGTWTCTAAT | 16S V4 (amplicon sequencing) |
| BFG5 | CCCCAAAAAGTCGCAAGGAC | pCBP54 detection |
| BFG6 | GACCGCTAACCAACACTCCA | pCBP54 detection |
| BFG7 | GTGGGACTGTTCGTCTTCGT | pAKP20 detection |
| BFG8 | CTGGGGATGATGTGCCTGTT | pAKP20 detection |
| BFG17 | CCTTGAACTGGTTTGTGCCG | pMRE25 detection |
| BFG18 | TGTGCTGATGAAGCGTCTGT | pMRE25 detection |
| BFG19 | AAAGCGTTTCGGTGTTCAGC | pBGP41 detection |
| BFG20 | AGAGAAGATGCGTATGGCGG | pBGP41 detection |
| BFG23 | TAGCCATCGTCTTCCATCGC | pPLE30.1 detection |
| BFG24 | CGCTCCTTCGTCGCTTATCT | pPLE30.1 detection |
| BFG25 | TGACGCTTTTGCTTGTGTCG | pPLE30.2 detection |
| BFG26 | TTGTTCGGTGATGTGAGCCA | pPLE30.2 detection |
| BFG27 | ACCACAACATCTACCCGCTC | pLP30.1 detection |
| BFG28 | CTTGGGCTCTCAGGTCACTC | pLP30.1 detection |
| BFG29 | GCAAGAGTGGACATACCCGA | pLP30.2 detection |
| BFG30 | AATGGTTGCTCTACTGGCGT | pLP30.2 detection |
| BFG31 | GTTCTCCGCCGATCTCTTGT | pMAP24.2 detection |
| BFG32 | TGGATTCGGTCTACGGCAAG | pMAP24.2 detection |
| BFG35 | GTCACGGTTGCTTTTCGGTG | pBPG46 detection |
| BFG36 | GTTTGCGTTGTCATGCCTCA | pBPG46 detection |

| | | |
|---|---|---|
| BFG39 | CATGCCGACCAGTGAAAGC | pPAP53 detection |
| BFG40 | GAAAAGTTCGCGTGGTCCTTC | pPAP53 detection |
| BFG41 | CACCACCAACGCTTTCCTTCT | pSHE10.1 detection |
| BFG42 | AGCTGCATCTGCGTCGATT | pSHE10.1 detection |
| BFG43 | AAGATCACCCCGAACAGGAAC | pSHE10.2 detection |
| BFG44 | CTTATGGTGGCACTCGCTTC | pSHE10.2 detection |
| BFG45 | GGCATCAGGTTTGGCTTCAG | pSPE13.4 detection |
| BFG46 | TCAAGGAGACGCTTTTGCTG | pSPE13.4 detection |
| BFG47 | CGAACCATCCTGAAACCCGA | pSPE13.1 detection |
| BFG48 | CATGGTCGGTGCCAGTAGAA | pSPE13.1 detection |
| BFG49 | GGGAACGAAGGTAAGCGACA | pSPE13.2 detection |
| BFG50 | CCAACGCTTTCCTTCTCAAG | pSPE13.2 detection |
| BFG51 | GCACGATCACGACGGTGTTG | pME13.1 detection |
| BFG52 | GGTGAATCAGTTCGTCTGCG | pME13.1 detection |
| BFG53 | GAGAGCGACTATCCCACCGA | pME13.2 detection |
| BFG54 | GGTGATGAAGTAGGCGAGCA | pME13.2 detection |
| BFG55 | CTCACGCAAGGGCTGTTTCT | pSPE13.3 detection |
| BFG56 | CGCCAGAAACCGCTCGTTA | pSPE13.3 detection |
| BFG57 | TTGGACTGCACATGAGGGTT | pLP73.1 detection |
| BFG58 | TGACAAATCGGCATCTCGGA | pLP73.1 detection |
| BFG59 | CAGACCGTACAAGAAGCCCT | pLP73.2 detection |
| BFG60 | CAGACCTCACGGTTGGTTGA | pLP73.2 detection |

† Phosphodiorate bond to prevent 3’ to 5’ exonuclease activity.

**Table S2: Taxonomic identification of metal-resistant isolates.** Isolate identities were determined through sequence alignment of the 16S rRNA gene. Either the entire gene was used, obtained from whole genome sequencing, or just the V3–V4 region of the 16S rRNA gene obtained by Sanger sequencing of 341F-805R amplicon.

| Isolate | Best type strain match (Accession number) | Identity (%) | GenBank whole genome accession |
|---|---|---|---|
| L2 | *Streptomyces nojiriensis* JCM 3382 (CP071139.1) | 100[a] | JBRECA000000000 |
| L3 | *Cupriavidus basilensis* DSM 11853 (CP062803.1) | 99.80[a] | JBRECI000000000 |
| L4 | *Paraburkholderia terrae* DSM 17804 (CP026111.1) | 99.74[a] | JBRECJ000000000 |
| L5 | *Streptomyces angustmyceticus* JCM 4053 (CP082945.1) | 99.54[a] | JBOBQK000000000 (*Streptomyces sirii*)[c] |
| L6 | *Streptomyces*[d] | 99.27[b] | |
| | *griseochromogenes* ATCC 14511(T) (MN686613.1) | | |
| | *hokutonensis* R1-NS-10 (NR_134197.1) | | |
| L7 | *Streptomyces chartreusis* ATCC 14922 (CP023689.1) | 98.95[a] | JBODSB000000000 |
| L8 | *Streptomyces galilaeus* ATCC 14969 (CP023703.1) | 99.08[a] | CP192594-6 |
| L9 | *Serratia* | 98.90[a] | CP192591-3 |
| | *liquefaciens* ATCC 27592 (NR_122057.1) | | |
| | *fonticola* DSM4576 (CP011254.1) | | |
| L10 | *Leifsonia poae* VKM Ac-1401 (DQ232613.2) | 99.39[a] | JBODSA000000000 |
| L11 | *Streptomyces*[d] | 99.75[b] | |
| | *bambusae* T110 (NR_146024.1) | | |
| | *kronopolitis* NEAU-ML8 (NR_153682.1) | | |
| L12 | *Streptomyces venezuelae* ATCC 10712 (CP029197.1) | 99.93[a] | CP192597 |
| L16 | *Ralstonia pickettii* FDAARGOS_1535 (CP084360.1) | 98.89[a] | JBNYXF000000000 |
| L17 | *Cupriavidus basilensis* DSM 11853 (CP062803.1) | 99.80[a] | JBRECE000000000 |
| L19 | *Pseudarthrobacter psychrotolerans* YJ56 (CP047898.1) | 98.62[a] | CP193935 |
| L23 | *Methylobacterium oryzae* CBMB20 (CP003811.1) | 99.80[a] | JBRECD000000000 |
| L25 | *Leifsonia xyli* subsp. *cynodontis* DSM 46306 (CP006734.1) | 98.88[a] | JBODRZ000000000 |
| L30 | *Enterobacter quasiroggenkampii* WCHECL1060 (KY979139.1) | 100[a] | CP192609-10 (*Enterobacter asburiae*)[c] |

| | | | |
|---|---|---|---|
| L33 | *Cupriavidus basilensis* DSM 11853 (CP062803.1) | 99.87[a] | JBRECB000000000 |
| L42 | *Cupriavidus basilensis* DSM 11853 (CP062803.1) | 100[b] | |
| P2 | *Dankookia rubra* WS-10 (NR_146664.1) | 98.44[a] | CP192581-2 |
| P3 | *Puia dinghuensis* 4GSH07 (NR_159128.1) | 97.20[a] | CP192583 |
| P4 | *Leifsonia poae* VKM Ac-1401 (DQ232613.2) | 99.46[a] | CP192580 |
| P6 | *Sinomonas atrocyanea* KCTC 3377 (CP014518.1) | 99.87[a] | JBRECG000000000 |
| P7 | *Mycolicibacterium psychrotolerans* JCM 13323 (AP022574.1) | 99.21[a] | CP192587-90 |
| P11 | *Streptomyces* | 100[b] | |
| | *albosporeus* subsp. *labilomyceticus* NBRC 15387 (NR_125445.1) | | |
| | *qinzhouensis* SSL-25 (CP042266.1) | | |
| P12 | *Leifsonia xyli* subsp. *cynodontis* DSM 46306 (CP006734.1) | 99.08[a] | CP192578 |
| P13 | *Leifsonia xyli* subsp. *cynodontis* DSM 46306 (CP006734.1) | 99.02[a] | CP192573 |
| P14 | *Methylobacterium* | 99.80[a] | JBOEFT000000000 |
| | *phyllosphaerae* CBMB27 (CP015367.1) | | |
| | *oryzae* CBMB27 (CP003811.1) | | |
| P15 | *Methylobacterium currus* PR1016A (CP028844.1) | 98.19[a] | |
| P17 | *Methylobacterium*[d] | 99.49[b] | |
| | *longum* 440 (NR_117045.1) | | |
| | *oryzae* CBMB20 (CP003811.1) | | |
| P19 | *Mucilaginibacter celer* HYN0043 (CP032869.1) | 98.09[a] | CP192586 |
| P20 | *Amnibacterium kyonggiense* KSL51201-037 (NR_116715.1) | 99.79[a] | JBREBW000000000 |
| P21 | *Flavobacterium zhairuonense* A5.7 (NR_165723.1) | 98.24[a] | CP192585 |
| P22 | *Caulobacter segnis* TK0059 (CP027850.1) | 99.53[a] | JBODRY000000000 |
| P23 | *Streptomyces*[d] | 99.75[b] | |
| | *gilvifuscus* T113 (NR_137389.1) | | |
| | *novaecaesareae* DSM 40358T (KF772673.1) | | |
| P24 | *Mesorhizobium atlanticum* CNPSo 3140 (MH128379.1) | 99.93[a] | JBREBX000000000 |
| P25 | *Mucilaginibacter celer* HYN0043 (CP032869.1) | 98.09[a] | CP193937 |
| P26 | *Pedobacter* sp. Marseille-Q2390 (LR797942.1) | 97.96[a] | CP192574 |

| | | | |
|---|---|---|---|
| P28 | *Ensifer adhaerens* Casida A (CP015880.2) | 100[a] | JBRECK000000000 |
| P29 | *Leifsonia*[d] | 99.74[b] | |
| | *aquatica* ATCC 14665 (MW228036.1) | | |
| | *xyli* subsp. *cynodontis* DSM 46306 (CP006734.1) | | |
| P30 | *Leifsonia xyli* subsp. *cynodontis* DSM 46306 (CP006734.1) | 98.88[a] | CP194644-6 |
| P31 | *Methylobacterium phyllosphaerae* CBMB27 (CP015367.1) | 98.85[a] | JBODSC000000000 |
| P32 | *Methylobacterium oryzae* CBMB20 (CP003811.1) | 99.80[a] | JBRECF000000000 |
| P33 | *Methylobacterium*[d] | 98.99[b] | |
| | *longum* 440 (NR_117045.1) | | |
| | *oryzae* CBMB20 (CP003811.1) | | |
| P34 | *Methylobacterium*[d] | 98.47[b] | |
| | *longum* 440 (NR_117045.1) | | |
| | *oryzae* CBMB20 (CP003811.1) | | |
| P35 | *Methylobacterium*[d] | 98.48[b] | |
| | *longum* 440 (NR_117045.1) | | |
| | *oryzae* CBMB20 (CP003811.1) | | |
| P36 | *Mitsuaria chitinivorans* HWN-4 (NR_175474.1) | 99.67[a] | CP192577 (*Roseateles chitinivorans*)[c] |
| P37 | *Mitsuaria chitinivorans* HWN-4 (NR_175474.1) | 99.60[a] | CP192575 (*Roseateles chitinivorans*)[c] |
| P38 | *Duganella lactea FT50W* (MN865818.2) | 98.93[a] | CP193936 |
| P39 | *Methylobacterium tardum* DSM 19566 (CP097484.1) | 99.80[a] | |
| P41 | *Bradyrhizobium guangxiense* CCBAU 53363 (NR_145894.1) | 99.80[a] | CP191781-2 |
| P42 | *Bradyrhizobium betae* PL7HG1 (CP044543.1) | 99.87[a] | CP192576 |
| P43 | *Bradyrhizobium betae* 39S1MB (CP029426.1) | 99.87[a] | JBRECH000000000 |
| P46 | *Bradyrhizobium guangxiense* CCBAU 53363 (CP022219.1) | 100[a] | JBREBY000000000 |
| P47 | *Mitsuaria chitinivorans* HWN-4 (NR_175474.1) | 99.60[a] | CP192584 (*Roseateles chitinivorans*)[c] |
| P48 | *Methylobacterium*[d] | 100[b] | |
| | *longum* 440 (NR_117045.1) | | |

| | | | |
|---|---|---|---|
| | *oryzae* CBMB20 (CP003811.1) | | |
| P49 | *Methylobacterium*[d] | 98.99[b] | |
| | *longum* 440 (NR_117045.1) | | |
| | *oryzae* CBMB20 (CP003811.1) | | |
| P51 | *Variovorax ureilyticus* UCM-2 (NR_169351.1) | 99.66[a] | CP192579 |
| P53 | *Pantoea agglomerans* FDAARGOS 1447 (CP077366.1) | 99.61[a] | |
| P54 | *Cupriavidus basilensis* DSM 11853 (CP062804.1) | 98.82[a] | JBNYXC000000000 |
| P66 | *Cupriavidus basilensis* DSM 11853 (CP062803.1) | 99.74[b] | |
| P71 | *Leifsonia xyli* subsp. *cynodontis* DSM 46306 (CP006734.1) | 91.69[a] | |
| P73 | *Leifsonia xyli* subsp. *cynodontis* DSM 46306 (CP006734.1) | 98.82[a] | CP194931-3 |
| E4 | *Streptomyces*[d] | 99.5[b] | |
| | *gilvifuscus* T113 (NR_137389.1) | | |
| | *novaecaesareae* DSM 40358T (KF772673.1) | | |
| E5 | *Streptomyces lasalocidi* ATCC 31180 (MK852399.1) | 99.28[a] | JBNYXE000000000 |
| E6 | *Streptomyces*[d] | 100[b] | |
| | *gilvifuscus* T113 (NR_137389.1) | | |
| | *novaecaesareae* DSM 40358T (KF772673.1) | | |
| E7 | *Streptomyces jiujiangensis* JXJ 0074 (NR_125706.1) | 98.87[a] | |
| E10 | *Sphingomonas hankookensis* ODN7 (NR_116570.1) | 99.45[a] | CP191783-8 |
| E12 | *Methylorubrum*[d] | 98.22[b] | |
| | *extorquens* TK 0001 (LT962688.1) | | |
| | *podarium* DSM 15083 (NR_112676.1) | | |
| E13 | *Sphingomonas panni* C52 (MW227641.1) | 99.86[a] | JBREBZ000000000 |
| E14 | *Mycolicibacterium aichiense* JCM 6376 (AP022561.1) | 99.6[a] | CP192571-2 |
| E15 | *Streptomyces*[d] | 99.25[b] | |
| | *gilvifuscus* T113 (NR_137389.1) | | |
| | *novaecaesareae* DSM 40358T (KF772673.1) | | |
| E16 | *Methylobacterium tardum* DSM 19566 (CP097484.1) | 99.80[a] | |
| E17 | *Yinghuangia aomiensis* M24DS4 (NR_112998.1) | 99.73[a] | JBRFIQ000000000 |
| E18 | *Streptomyces candidus* JCM 4629 (MT760586.1) | 99.74[b] | |

| | | | |
|---|---|---|---|
| E19 | *Bacillus cereus* subsp.[d] | 99.51[b] | |
| | *cereus* CCM 2010 (MT421928.1) | | |
| | *mycoides* NBRC 101228 (MF347936.1) | | |
| E20 | *Bradyrhizobium oligotrophicum* S58 (NR_102489.2) | 99.80[a] | CP192619-20 |
| E21 | *Methylobacterium*[d] | 98.74[b] | |
| | *longum* 440 (NR_117045.1) | | |
| | *oryzae* CBMB20 (CP003811.1) | | |
| E22 | *Rhodococcus pseudokoreensis* R79 (CP070619.1) | 97.83[a] | CP194064 *(Prescottella equi)*[c] |
| E23 | *Streptomyces*[d] | 99.75[b] | |
| | *gilvifuscus* T113 (NR_137389.1) | | |
| | *novaecaesareae* DSM 40358T (KF772673.1) | | |
| E24 | *Methylobacterium* | 98.23[b] | |
| | *ajmalii* IF7SW-B2 (MW532474.1) | | |
| | *indicum* SE2.11 (NR_135210.1) | | |
| E25 | *Methylobacterium radiotolerans* JCM 2831 (CP001001.1) | 99.93[a] | JBNYXH000000000 |
| E27 | *Methylorubrum*[d] | 99.22[b] | |
| | *extorquens* TK 0001 (LT962688.1) | | |
| | *podarium* DSM 15083 (NR_112676.1) | | |
| E29 | *Methylorubrum*[d] | 99.48[b] | |
| | *extorquens* TK 0001 (LT962688.1) | | |
| | *podarium* DSM 15083 (NR_112676.1) | | |
| E30 | *Pigmentiphaga litoralis* JSM 061001 (NR_044530.1) | 99.80[a] | CP184479-80 |
| E31 | *Sphingomonas hankookensis* ODN7 (NR_116570.1) | 99.45[a] | JBNYXD000000000 |
| E32 | *Methylobacterium radiotolerans* JCM 2831 (NR_074244.1) | 98.74[b] | |
| E40 | *Cupriavidus basilensis* DSM 11853 (CP062803.1) | 99.87[a] | JBRECC000000000 |

a- Type strain alignment with entire 16S rRNA gene
b- Type strain alignment with V3-V4 region of 16S rRNA gene
c- Renamed by NCBI based on ANI
d- Too many equal matches to list, two listed for brevity

**Table S3: Summary statistics for Oxford Nanopore sequencing reads of pooled bacterial whole genome extracts.** Filtered reads refers to reads with lengths exceeding 1 kb.

| Pooled sample | No. of DNA extracts | Total reads | Total Mb | Filtered reads (%) | Filtered bp (%) | Filtered mean length (bp) | Filtered quality | Longest read (kb) | Contigs | Average coverage/contig | Contigs per DNA extract |
|---|---|---|---|---|---|---|---|---|---|---|---|
| S1 | 2 | 749,822 | 1853 | 42.94 | 88.56 | 5096 | 12.1 | 353 | 17 | 38.24 | 9.5 |
| S2 | 3 | 192,514 | 1011 | 61.85 | 96.45 | 8193 | 12.3 | 489 | 107 | 7.2 | 35.67 |
| S3 | 8 | 1,287,314 | 6605 | 69.28 | 96.62 | 7155.2 | 13.5 | 162 | 89 | 35.53 | 8.9 |
| S4 | 8 | 1,256,408 | 2671 | 48.44 | 86.8 | 3808.4 | 11 | 457 | 518 | 8.05 | 64.75 |
| S5 | 8 | 423,243 | 1529 | 58.43 | 93.92 | 5804.8 | 11.1 | 314 | 214 | 9.71 | 26.75 |
| S6 | 8 | 372,488 | 1259 | 52.7 | 92.3 | 5917.7 | 11.1 | 116 | 411 | 6.1 | 51.38 |
| S7 | 8 | 286,201 | 1188 | 55.31 | 94.6 | 7098.8 | 11.2 | 103 | 183 | 5.42 | 22.88 |
| S8 | 8 | 461,954 | 1688 | 53.87 | 93.6 | 6350.6 | 11.9 | 110 | 234 | 11.71 | 29.25 |
| S9 | 5 | 336,874 | 1647 | 65.5 | 96.4 | 7197 | 12 | 1097 | 270 | 9.48 | 54 |
| S10 | 4 | 649,538 | 2429 | 52.61 | 93.64 | 6657.5 | 11.7 | 100 | 72 | 55.11 | 18 |
| S11 | 3 | 142,377 | 708.8 | 73.26 | 97.27 | 6609.2 | 13.1 | 455 | 27 | 17.19 | 9 |
| S12 | 4 | 148,915 | 894.9 | 74.35 | 97.87 | 7911.3 | 13 | 62 | 201 | 18.83 | 50.25 |

**Table S4: Plasmids detected by whole genome sequencing and their features.** Plasmids were identified as circularised sequences, <2 Mb, sequence coverage ≥5, and not carrying rRNA genes. Predicted host is the isolate that contributed DNA to the pooled sample that belongs to the same taxonomic family as the BLASTN alignment sequence of the contig. Host prediction was confirmed by PCR for plasmids containing metal or antibiotic resistance genes. Plasmid mobility type (MOB) predicted by MOBscan [41], based on relaxase gene identity. Plasmids without relaxase genes are classified as not applicable (NA), whereas plasmids with relaxase genes that did not belong to a known group are classified as novel.

| Pooled Sample | Contig | Size (bp) | Sequence coverage | MOB type | BLASTN plasmid sequence alignment (accession no.) | Nucleotide identity (%) | Query coverage (%) | Predicted host | Notes |
|---|---|---|---|---|---|---|---|---|---|
| S2 | 107 | 159,617 | 33 | Novel | *Agrobacterium larrymoorei* CFBP5473 pTiCFBP5473 (CP039694.1) | 91.2 | 2 | *Ensifer adhaerens* P28 | |
| S2 | 109 | 171,550 | 7 | NA | *Cupriavidus basilensis* DSM 11853 pRK1-3 (CP062807.1) | 99.29 | 15 | *C. basilensis* P54 | PCR confirmed to host, named pCBP54 |
| S3 | 3 | 110,749 | 74 | NA | *Streptomyces violaceoruber* CGMCC 4.1801 (CP137734.1) | 84.67 | 21 | *Streptomyces galilaeus* L8 | |
| S3 | 76 | 24,468 | 32 | $MOB_F$ | *Mycolicibacterium crocinum* JCM 16369 plasmid unnamed2 (CP103314.1) | 94.63 | 26 | *Mycolicibacterium psychrotolerans* P7 | |
| S3 | 99 | 351,110 | 70 | $MOB_F$ | *M. crocinum* JCM 16369 plasmid unnamed (CP092363.2) | 91.5 | 3 | *M. psychrotolerans* P7 | |
| S3 | 101 | 55,044 | 33 | NA | *Methylobacterium phyllosphaerae* CBMB27 CBMB27-p1 (CP015368.1) | 87.44 | 1 | *M. phyllosphaerae* P14 | PCR confirmed to host |

| | | | | | | | | |
|---|---|---|---|---|---|---|---|---|
| S3 | 102 | 162,624 | 350 | Novel | *Serratia nevei* LMG 31536 pSNEVa (CP149941.1) | 84.63 | 11 | *Serratia fonticola* L9 | |
| S3 | 103 | 51,845 | 144 | $MOB_P$ | *Clavibacter phaseoli* LPPA 982 pCP (CP040787.1) | 83.53 | 2 | *Amnibacterium kyonggiense* P20 | PCR confirmed to host, named pAKP20 |
| S3 | 104 | 27,183 | 26 | NA | *Methylobacterium currus* PR1016A plasmid unnamed1 (CP028845.1) | 83.81 | 5 | *M. phyllosphaerae* P14 | |
| S4 | 239 | 5554 | 8 | NA | *Herbaspirillum hiltneri* N3 (CP011409.1) | 77.13 | 44 | *Duganella* sp. P38 | |
| S4 | 550 | 46,586 | 11 | $MOB_F$ | *Methylobacterium currus* PR1016A plasmid unnamed1 (CP028845.1) | 88.61 | 23 | *Methylobacterium radiotolerans* E25 | PCR confirmed to host, named pMRE25 |
| S4 | 551 | 33,127 | 17 | $MOB_F$ | *M. radiotolerans* JCM 2831 pMRAD03 (CP001004.1) | 96.12 | 31 | *M. radiotolerans* E25 | |
| S4 | 556 | 139,618 | 23 | Novel | *Afipia carboxidovorans* OM5 pOC167 (CP002828.1) | 79.77 | 17 | *Bradyrhizobium betae* P43 | |
| S5 | 54 | 248,638 | 25 | $MOB_P$ | *Bradyrhizobium elkanii* USDA 76 pUSDA76 (CP066357.1) | 83.15 | 14 | *Bradyrhizobium guangxiense* P41 | PCR confirmed to host, named pBGP41 |
| S5 | 142 | 81,805 | 55 | $MOB_P$ | *Sinomonas atrocyanea* KCTC 3377 pSA01 (CP014519.1) | 89.79 | 12 | *S. atrocyanea* P6 | |
| S5 | 143 | 152,219 | 45 | $MOB_P$ | *S. atrocyanea* KCTC 3377 pSA01 (CP014519.1) | 95.23 | 18 | *S. atrocyanea* P6 | |

| | | | | | | | | | |
|---|---|---|---|---|---|---|---|---|---|
| S5 | 165 | 100,066 | 52 | $MOB_P$ | *S. atrocyanea* KCTC 3377 pSA01 (CP014519.1) | 97.43 | 29 | *S. atrocyanea* P6 | PCR confirmed to different host, *Pigmentiphaga litoralis* E30, named pPLE30.1 |
| S5 | 220 | 49,732 | 19 | NA | *Methylobacterium durans* 17SD2-17 (CP029550.1) | 83.05 | 21 | *Methylobacterium oryzae* P32 | |
| S5 | 225 | 199,488 | 57 | NA | *Burkholderia contaminans* LMG 23361 plasmid unnamed2 (CP090644.1) | 74.53 | 2 | *P. litoralis* E30 | PCR confirmed to host, named pPLE30.2[†] |
| S5 | 230 | 39,123 | 15 | NA | *Methylobacterium tardum* DSM 19566 (CP097484.1) | 91.29 | 16 | *M. oryzae* P32 | |
| S5 | 232 | 22,202 | 8 | NA | *M. radiotolerans* JCM 2831 (CP001001.1) | 80.64 | 4 | *M. oryzae* P32 | |
| S5 | 233 | 10,295 | 12 | NA | No similarity | | | Unknown | |
| S6 | 94 | 264,303 | 13 | NA | *C. basilensis* DSM 11853 (CP062803.1) | 93.11 | 13 | *C. basilensis* L17 | |
| S6 | 178 | 49,884 | 132 | NA | *Bradyrhizobium cosmicum* 58S1 (CP041656.1) | 76.62 | 4 | *Bradyrhizobium oligotrophicum* E20 | |
| S7 | 178 | 57,599 | 92 | $MOB_P$ | *Leucobacter triazinivorans* JW-1 (CP035806.1) | 82.26 | 6 | *Leifsonia* sp. P30 | PCR confirmed to host, named pLP30.1 |
| S7 | 184 | 128,795 | 105 | NA | *Clavibacter michiganensis* subsp. *tessellarius* ATCC 33566 pCT2 (CP040790.1) | 89.7 | 1 | *Leifsonia* sp. P30 | PCR confirmed to host, named pLP30.2 |
| S8 | 153 | 202,928 | 32 | $MOB_F$ | *Sphingomonas paucimobilis* FDAARGOS_908 plasmid unnamed1 (CP065669.1) | 99.94 | 54 | *Sphingomonas hankookensis* E31 | PCR confirmed to host, identical to S9 contig 27 |

| | | | | | | | | | |
|---|---|---|---|---|---|---|---|---|---|
| S8 | 201 | 30,285 | 16 | Novel | *M. oryzae* CBMB20 pMOC2 (JX627581.1) | 90.32 | 23 | *M. tardum* E16 | |
| S8 | 219 | 381,974 | 8 | Novel | *Mesorhizobium jarvisii* ATCC 33669 pMJ33669a (CP033508.1) | 86.25 | 2 | *Mesorhizobium atlanticum* P24 | PCR confirmed to host, named pMAP24.2 |
| S8 | 220 | 36,211 | 31 | $MOB_F$ | *M. phyllosphaerae* y76CBMB27 CBMB27-p2 (CP015369.1) | 91.65 | 3 | *M. tardum* E16 | |
| S8 | 224 | 180,240 | 43 | $MOB_P$ | *Bradyrhizobium septentrionale* 1S1 pBs1S1a (CP088284.1) | 90.58 | 23 | *B. guangxiense* P46 | PCR confirmed to host, named pBGP46 |
| S8 | 233 | 125,010 | 132 | $MOB_F$ | *Sphingomonas sanxanigenens* DSM 19645 = NX02 pNXO2 (CP011450.1) | 77.82 | 11 | *S. hankookensis* E31 | Identical to S9 contig 266 |
| S8 | 236 | 89,102 | 71 | NA | *S. paucimobilis* FDAARGOS_908 plasmid unnamed1(CP065669.1) | 99.95 | 7 | *S. hankookensis* E31 | |
| S8 | 246 | 218,256 | 49 | $MOB_F$ | *M. crocinum* JCM 16369 plasmid unnamed (CP092363.2) | 91.69 | 2 | *Mycolicibacterium aichiense* E14 | |
| S8 | 248 | 22,267 | 18 | NA | *M. tardum* DSM 19566 pMt-1 (CP097485.1) | 86.65 | 5 | *M. tardum* E16 | |
| S9 | 27 | 202,926 | 149 | $MOB_F$ | *S. paucimobilis* FDAARGOS_908 plasmid unnamed1 (CP065669.1) | 99.96 | 54 | *S. hankookensis* E10 | PCR confirmed to host, identical to S8 contig 153, named pSHE10.1 |
| S9 | 137 | 239,968 | 115 | $MOB_F$ | *Sphingomonas aliaeris* DH-S5 plasmid unnamed1 (CP061036.1) | 79.61 | 11 | *S. hankookensis* E10 | Identical to S10 contig 47 |
| S9 | 138 | 64,252 | 173 | $MOB_P$ | *Sphingomonas insulae* KCTC 12872 plasmid unnamed1 (CP048419.1) | 88.63 | 44 | *S. hankookensis* E10 | PCR confirmed to host, named pSHE10.2 |

| | | | | | | | | | |
|---|---|---|---|---|---|---|---|---|---|
| S9 | 266 | 125,003 | 385 | $MOB_F$ | *S. sanxanigenens* DSM 19645 = NX02 pNXO2 (CP011450.1) | 77.82 | 11 | *S. hankookensis* E10 | |
| S9 | 269 | 89,090 | 273 | NA | *S. paucimobilis* FDAARGOS_908 plasmid unnamed1 (CP065669.1) | 99.92 | 7 | *S. hankookensis* E10 | |
| S9 | 272 | 1,892,539 | 44 | NA | *Ideonella dechloratans* CCUG 30977 plasmid unnamed1 (CP088082.1) | 84.33 | 1 | Unknown | PCR confirmed to *Pantoea agglomerans* P53, named pPAP53 |
| S10 | 2 | 206,010 | 294 | NA | *S. paucimobilis* FDAARGOS_908 plasmid unnamed1 (CP065669.1) | 99.72 | 29 | *Sphingomonas panni* E13 | PCR confirmed to host, named pSPE13.1 |
| S10 | 3 | 69,420 | 531 | $MOB_F$ | *S. paucimobilis* FDAARGOS_908 plasmid unnamed1 (CP065669.1) | 99.85 | 26 | *S. panni* E13 | PCR confirmed to host, named pSPE13.2 |
| S10 | 4 | 38,227 | 348 | NA | *S. paucimobilis* FDAARGOS_908 plasmid unnamed1 (CP065669.1) | 99.92 | 33 | *S. panni* E13 | PCR confirmed to host, named pSPE13.3 |
| S10 | 5 | 241,771 | 213 | $MOB_F$ | *Sphingobium cloacae* JCM 10874 pSCLO_2 (AP017656.1) | 98.71 | 20 | *S. panni* E13 | PCR confirmed to host, named pSPE13.4 |
| S10 | 47 | 229,691 | 484 | $MOB_F$ | *S. paucimobilis* FDAARGOS_908 plasmid unnamed1 (CP065669.1) | 90.50 | 4 | *S. panni* E13 | Identical to S9 contig 137 |
| S10 | 58 | 28,794 | 36 | NA | *M. oryzae* CBMB20 (CP003811.1) | 91.73 | 12 | *Methylobacterium* sp. P31 | |
| S10 | 65 | 73,142 | 365 | NA | *Sphingomonas radiodurans* S9-5 (CP086594.1) | 83.29 | 10 | *S. panni* E13 | |
| S10 | 67 | 102,466 | 653 | $MOB_F$ | *Sphingomonas wittichii* RW1 pSWIT01 (CP000700.1) | 75.67 | 9 | *S. panni* E13 | |

| | | | | | | | | | |
|---|---|---|---|---|---|---|---|---|---|
| S10 | 71 | 64,947 | 14 | Novel | *M. currus* PR1016A plasmid unnamed1 (CP028845.1) | 99.84 | 13 | *Methylobacterium* sp. P31 | Not confirmed to host, named pME13.1 |
| S10 | 72 | 45,939 | 8 | $MOB_F$ | *Methylobacterium populi* BJ001 (CP001029.1) | 99.86 | 13 | *Methylobacterium* sp. P31 | |
| S10 | 73 | 18,424 | 5 | NA | *M. currus* PR1016A plasmid unnamed1 (CP028845.1) | 88.05 | 3 | *Methylobacterium* sp. P31 | |
| S10 | 74 | 30,442 | 27 | NA | *M. radiotolerans* JCM 2831 pMRAD01 (CP001002.1) | 91.52 | 5 | *Methylobacterium* sp. P31 | |
| S10 | 76 | 48,756 | 7 | $MOB_F$ | *M. currus* PR1016A plasmid unnamed1 (CP028845.1) | 88.05 | 3 | *Methylobacterium* sp. P31 | Not confirmed to host, named pME13.2 |
| S11 | 4 | 68,729 | 19 | $MOB_P$ | *Microbacterium pygmaeum* DSM 23142 (LT629692.1) | 77.25 | 5 | *Leifsonia* sp. P73 | PCR confirmed to host, named pLP73.1 |
| S11 | 5 | 69,333 | 15 | $MOB_P$ | *M. pygmaeum* DSM 23142 (LT629692.1) | 76.93 | 5 | *Leifsonia* sp. P73 | PCR confirmed to host, named pLP73.2 |
| S11 | 34 | 8646 | 110 | NA | *Enterobacter hormaechei* subsp. *oharae* FDAARGOS_1533 plasmid unnamed2 (CP083615.1) | 88.33 | 13 | *Enterobacter quasiroggenkampii* L30 | |
| S12 | 233 | 4323 | 92 | NA | *E. hormaechei* subsp. *oharae* FDAARGOS_1533 plasmid unnamed2 (CP083615.1) | 88.51 | 13 | Unknown | |
| S12 | 235 | 292,707 | 25 | $MOB_H$ | *Cupriavidus necator* N-1 pBB1 (CP002879.1) | 74.93 | 5 | *C. basilensis* L33 | |

† Described in [57]

**Table S5: Inhibition zones from Calibrated Dichotomous Susceptibility disc diffusion test.** Mean annular radii of inhibition zones (mm) measured 24–48 hours post inoculation (*n* = 3, standard deviation). Inhibition zones representing resistance (<6 mm) are in bold.

| Antibiotic class | Aminoglycosides | | | | | | β-lactams | | | | Quinolones | | Folate inhibitors | | Tetracyclines | | Phenicols | Macrolides |
|---|---|---|---|---|---|---|---|---|---|---|---|---|---|---|---|---|---|---|
| Strain | Am | Gen | Kan | Neo | Spt | Str | Amp | Cro | Ctx | Ipm | Cip | Nal | Sul | Tmp | Dox | Tet | Chl | Ery |
| *Amnibacterium kyonggiense* P20 | >20 | >20 | >20 | >20 | >20 | >20 | >20 | >20 | >20 | >20 | >20 | >20 | >20 | >20 | >20 | >20 | >20 | >20 |
| *Bradyrhizobium guangxiense* P41 | >20 | >20 | >20 | >20 | >20 | >20 | 10.5 ± 0.7 | >20 | >20 | >20 | 7.7 ± 0.6 | 11 ± 2.6 | >20 | >20 | >20 | >20 | 19 ± 1.4 | >20 |
| *Bradyrhizobium guangxiense* P46 | 14.5 ± 0.5 | 14.5 ± 3.5 | **5 ± 1.5** | 18.2 ± 1.3 | >20 | 17.5 ± 1.8 | **1.2 ± 0.8** | 12.8 ± 1.8 | 16 ± 2.6 | 17.4 ± 1.2 | 7 ± 0.9 | **0.3 ± 0.6** | 8 ± 1.4 | **0.5 ± 0.5** | **4 ± 2.3** | **4.7 ± 1.9** | **1.1 ± 1.5** | 8.5 ± 0.7 |
| *Cupriavidus basilensis* P54 | 11.3 ± 0.6 | 10.7 ± 1.5 | 11.7 ± 1.5 | 9.3 ± 1.2 | 8 ± 1.7 | 8.5 ± 0.5 | 6.3 ± 0.6 | 15.7 ± 1.5 | 18.3 ± 1.5 | >20 | 14.3 ± 0.6 | **5.7 ± 0.6** | 15.3 ± 2.1 | **4 ± 1.0** | 11.5 ± 0.9 | 10.3 ± 0.6 | **5.7 ± 0.6** | **2.7 ± 0.3** |
| *Leifsonia* sp. P30 | >20 | >20 | >20 | >20 | >20 | >20 | >20 | >20 | >20 | >20 | 16 ± 1.4 | **2.5 ± 1.4** | >20 | >20 | >20 | >20 | >20 | >20 |
| *Leifsonia* sp. P73 | 14.8 ± 2.8 | 9.2 ± 2.0 | 15 ± 2.0 | 9.2 ± 2.0 | >20 | 15 ± 1.0 | 14.5 ± 0.7 | 19 ± 1.0 | 18.7 ± 1.5 | >20 | 15.5 ± 0.7 | **0.5 ± 0.7** | >20 | 16.5 ± 2.1 | >20 | 17.8 ± 1.3 | 14 ± 1.4 | >20 |
| *Mesorhizobium atlanticum* P24 | >20 | >20 | >20 | >20 | >20 | >20 | >20 | >20 | >20 | >20 | 18.3 ± 0.7 | **3 ± 1.4** | >20 | >20 | >20 | >20 | >20 | >20 |

| | | | | | | | | | | | | | | | | | | |
|---|---|---|---|---|---|---|---|---|---|---|---|---|---|---|---|---|---|---|
| *Methylobacterium radiotolerans* E25 | 12.5 ± 1.4 | 8.3 ± 1.1 | 17.3 ± 2.5 | 6.8 ± 1.1 | 20 ± 0.0 | 14.3 ± 1.8 | **0 ± 0.0** | 16.5 ± 2.1 | 9 ± 0.0 | >20 | 11 ± 2.8 | **0 ± 0.0** | **0 ± 0.0** | **0 ± 0.0** | 17.8 ± 3.2 | 17.5 ± 0.7 | **3.5 ± 2.1** | 17.5 ± 3.5 |
| *Pantoea agglomerans* P53 | 11.8 ± 1.8 | 7.8 ± 1.1 | 13.5 ± 4.9 | 8.3 ± 1.1 | >20 | 13.5 ± 3.5 | 18 ± 2.8 | 15 ± 0.7 | 19.5 ± 0.7 | >20 | 17.8 ± 0.4 | 12.5 ± 2.1 | >20 | 15.7 ± 5.1 | >20 | 19.5 ± 0.5 | 15 ± 0.0 | 17.5 ± 0.7 |
| *Pigmentiphaga litoralis* E30 | 13 ± 1.3 | 7.7 ± 0.8 | 12.3 ± 1.5 | **5.3 ± 0.3** | 17.7 ± 2.1 | 11.7 ± 1. | **0.3 ± 0.3** | 7.5 ± 2.6 | 6.7 ± 1.6 | 16 ± 1.0 | 18 ± 0 | 16 ± 1.3 | >20 | 13.3 ± 2.5 | 18.2 ± 0.8 | 14.2 ± 1.2 | 9 ± 2.2 | 7.7 ± 2.1 |
| *Sphingomonas hankookensis* E10 | 17.3 ± 1.5 | 16.2 ± 0.3 | 19 ± 1.0 | 13.7 ± 1.2 | 8.3 ± 1.0 | **1.2 ± 1.6** | 15.7 ± 3.1 | 16.3 ± 1.2 | 17 ± 0.0 | >20 | 9 ± 1.0 | **1 ± 1.0** | ND | 10 ± 1.7 | 18.3 ± 0.6 | 16 ± 0.0 | 12.2 ± 2.0 | 14.7 ± 1.2 |

Amk = amikacin, Amp = ampicillin, Chl = chloramphenicol, Cip = ciprofloxacin, Cro = ceftriaxone, Ctx = cefotaxime, Dox = doxycycline, Ery = erythromycin, Gen = gentamicin, Ipm = imipenem, Kan = kanamycin, Nal = nalidixic acid, Neo = neomycin, Spt = spectinomycin, Str = streptomycin, Sul = compound sulfonamides, Tet = tetracycline, Tmp = trimethoprim, ND = inhibition zone not distinct.

**Table S6: Putative transposon and insertion sequence (IS) regions identified in resistance plasmids.** IS predicted by ISFinder [44]. Position of IS in the plasmid is indicated along with the presence of left and right inverted repeats (IRL/IRR), difference in GC content (%) between the putative IS region and the plasmid backbone, in addition to other notable features detected within these regions. Novel IR detected with Inverted Repeat Finder [45].

| Plasmid | Nucleotide position | IS/transposon | IRL/IRR | ΔGC% of IS/transposon from total plasmid | Notable features |
|---|---|---|---|---|---|
| pMRE25 | 13,781–15,127 | ISM*tsp5* | IRL | +7 | |
| pBGP41 | 157,279–158,723 | IS*Bj6_B* | IRL | -3 | |
| pMAP24.2 | 30,227–30,426 | IS*Mlo4* | IRR | -2 | |
| pSHE10.1 | 14,601–16,155 | Tn*5393* | IRR | +1 | Left flank adjacent to β-lactamase gene |
| | 80,051–80,810 | IS*Kpn12* | IRL | -1 | |
| | 96,089–99,022 | IS*Gbe1* | IRL, IRR | 0 | |
| | 120,070–121,338 | IS*Gbe1* | IRL, IRR | 0 | Right flank adjacent to copper oxidase gene |
| | 161,923–164,473 | Tn*5393* | IRR | +2 | |
| | 173,556–176,163 | IS*Gbe1* | IRL | +5 | |
| pPAP53 | 199,252–200,899 | IS*Pa56* | IRL, IRR | -1 | |
| | 611,351–612,725 | IS*Pa56* | IRL, IRR | -1 | |
| pSPE13.1 | 204,273–205,826 | Tn*5393* | IRR | -2 | Left flank adjacent to β-lactamase gene |
| | 84,990–85,562 | IS*Sphsp7* | IRR | -2 | |
| | 99,171–101,513 | IS*Sphsp7* | IRL | 0 | |
| | 105,507–119,770 | IS*110*, IS*Rm31* | IRL, IRR | -1 | Novel IR |
| | 107,607–119,234 | IS*110*, IS*Rm31* | IRL, IRR | -1 | Novel IR |
| pSPE13.2 | 10,392–11,659 | IS*Gbe1* | IRL, IRR | +1 | |

| | | | | | |
|---|---|---|---|---|---|
| | 46,611–49,842 | IS*MPo10* | IRR | +4 | Left flank adjacent to β-lactamase gene |
| pSPE13.3 | 3–8172 | IS*481* | IRL, IRR | +3 | Novel IR |
| pSPE13.4 | 17,534–18,802 | IS*Gbe1* | IRL, IRR | 0 | |
| | 25,995–27,264 | IS*Gbe1* | IRL, IRR | 0 | |
| | 48,301–49,569 | IS*Gbe1* | IRL, IRR | 0 | Right flank adjacent to copper oxidase gene |
| | 86,223–90,302 | Tn*5393* | IRR | 0 | |
| | 99,381–100,649 | IS*Gbe1* | IRL, IRR | 0 | |
| | 170,829–173,760 | IS*Gbe1* | IRL, IRR | 0 | |
| | 223,480–231,647 | IS*481* | IRL, IRR | +3 | Novel IR |
| pME13.1 | 16–3879 | IS*Mex22* | IRL, IRR | +7 | |
| pME13.2 | 17,115–18,470 | IS*Mno26* | IRL, IRR | +3 | |
| | 24,580–26,332 | IS*Mex10* | IRL, IRR | +7 | |
| | 26,331–27,608 | IS*Mex5* | IRR | +2 | |
| | 28,467–28,755 | IS*Mex10* | IRL | +5 | |

**Table S7: Summary statistics for Oxford Nanopore sequencing reads from each plasmid metagenome.** Sequence collections from sediment samples (P, E) are abbreviated: 0 = original sediment, 1 = sediment amended with 2 mM Cd and Cr for 4 weeks, 2 = sediment amended with 20 mM Cd and Cr for 4 weeks, neg = *Escherichia coli* TOP10 inoculated into sterile sand. Filtered reads were greater than 1 kb in length and in the top 90% of quality scores. Retained contigs lacked rRNA genes, had coverage ≥5, and did not match to contigs in the negative control. NA = not applicable

| Sample | Total reads | Total kb | Filtered reads (%) | Filtered bp (%) | Filtered mean length (kb) | Filtered read quality | Longest read (kb) | Contigs | Retained contigs (%) | Mean retained contig length (kb) | Retained contigs >20 coverage (%) | Retained circularised contigs (%) |
|---|---|---|---|---|---|---|---|---|---|---|---|---|
| P0 | 98,836 | 469,047 | 69.21 | 90 | 6.17 | 14.7 | 92 | 231 | 86.6 | 5.45 | 60 | 24 |
| P1 | 61,286 | 316,678 | 47.47 | 90 | 9.8 | 17 | 160.6 | 577 | 59.4 | 23.7 | 12.2 | 12.8 |
| P2 | 37,925 | 219,058 | 49.73 | 90 | 10.45 | 16.6 | 168 | 352 | 69.9 | 32.49 | 6.91 | 10.6 |
| E0 | 101,115 | 476,949 | 48.44 | 90 | 8.87 | 17.2 | 226 | 1017 | 72.3 | 25.57 | 12.4 | 17.6 |
| E1 | 95,097 | 425,811 | 52.63 | 90 | 7.66 | 16.8 | 161 | 327 | 45.9 | 6.24 | 13.8 | 17.1 |
| E2 | 82,362 | 464,372 | 54.76 | 90 | 9.27 | 16.4 | 209 | 515 | 72 | 28.17 | 12.7 | 26.4 |
| Neg | 79,652 | 440,192 | 67.44 | 90 | 7.37 | 16.9 | 116.5 | 816 | NA | NA | NA | NA |

**Table S8: Metagenomic plasmid contigs that align with genes and replicons in plasmid-specific databases.** Alignments were conducted using PlasmidFinder for plasmid replicons [40], MOBscan for plasmid type based on relaxase genes [41], and oriTfinder for mobilisation apparatus genes [42].

| PlasmidFinder replicon | No. of contigs carrying gene/replicon alignment |
|---|---|
| colRNAI_1 | 5 |
| MOBscan gene | |
| $MOB_F$ | 4 |
| $MOB_H$ | 2 |
| $MOB_Q$ | 9 |
| $MOB_P$ | 4 |
| $MOB_V$ | 4 |
| OriTfinder gene | |
| oriT | 1 |
| Relaxase | 3 |
| Type 4 coupling protein | 2 |

**Table S9: Plasmid contigs from sediment metagenomes exhibiting features of co-selection.** Co-selection models are classified as co-resistance, defined as the presence of both a metal resistance gene (MRG) and an antibiotic resistance gene (ARG); or cross-resistance, defined as the presence of a gene conferring resistance to both metal and antibiotic. Gene homologues are indicated when alignment scores are lower than those for specifically names genes.

| Contig | MRGs | ARGs | Co-selection model |
|---|---|---|---|
| E0_54 | *arsCBH* | *acrAD* | Co-resistance |
| E0_210 | *arsA*-homologue, *macAB*-homologue | *macAB*-homologue | Co-, cross-resistance |
| E0_279 | *chrR*-homologue | *emrAB*-homologue | Co-resistance |
| E0_358 | *copA*-homologue | *arr*-homologue | Co-resistance |
| E0_374 | *mdtABC* | *mdtABC* | Cross-resistance |
| E0_438 | *mdtA*-homologue | *mcrA*, *acrB*, *mdtA*-homologue, $bla_{\text{OXA-42}}$, $bla_{\text{PEN-J}}$ | Co-, cross-resistance |
| E0_486 | *actP*, *czcA*-homologue, *mdtA*-homologue | *mdtA*-homologue | Co-, cross-resistance |
| E0_1328 | *cmeB* | *cmeB* | Cross-resistance |
| P1_41 | *cadA* | *arnTFBCA* | Co-resistance |
| P1_589 | *silP, gesB* | *gesB*, *bepG*-homologue | Co-, cross-resistance |
| P2_24 | *cadA, czcICBARS* | *arnAD* | Co-resistance |
| P2_178 | *mdtABC* | *mdtABC* | Cross-resistance |
| P2_447 | *actP* | *rosB*-homologue | Co-resistance |

A

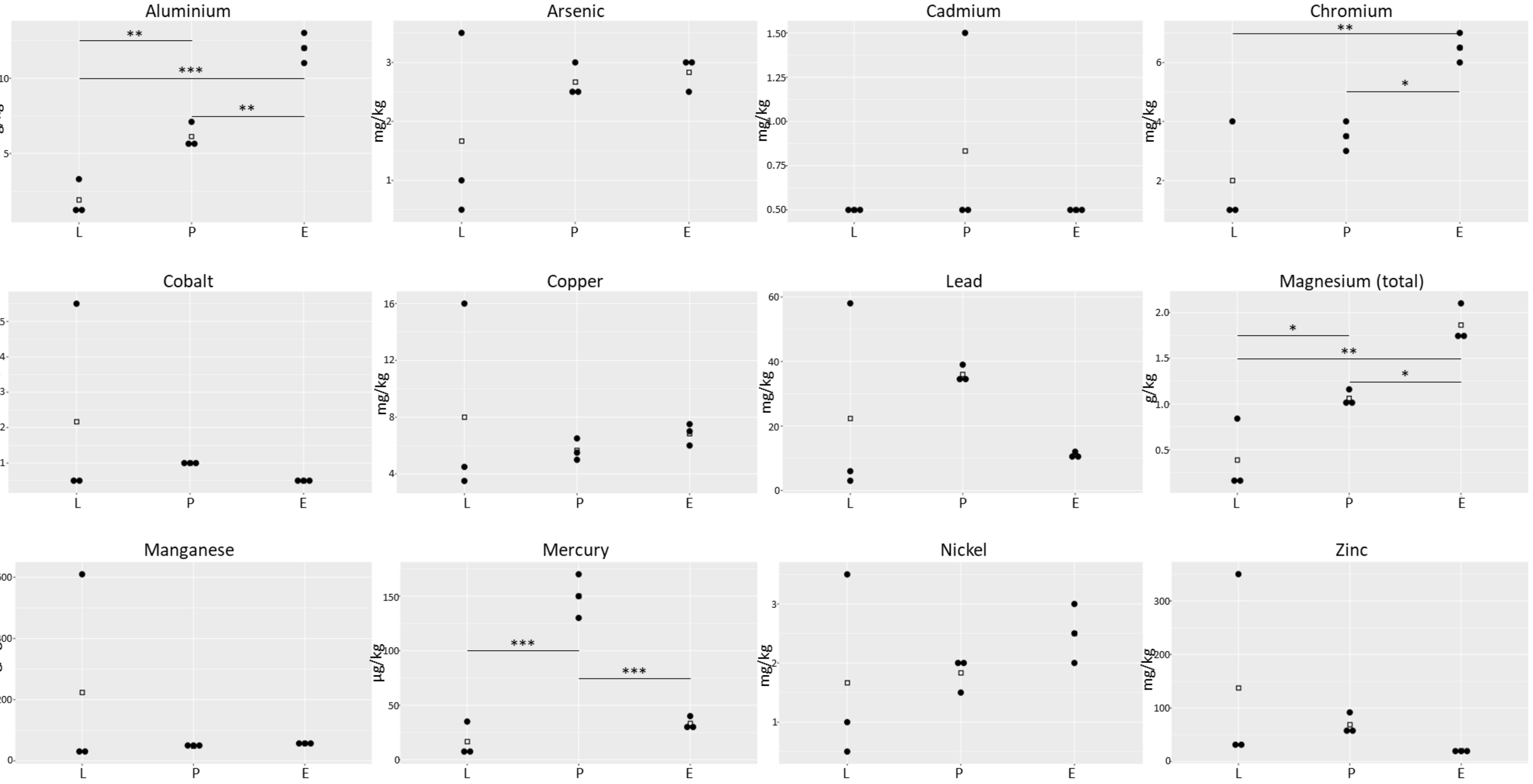
Aluminium
g/kg
Arsenic
mg/kg
Cadmium
mg/kg
Chromium
mg/kg
Cobalt
mg/kg
Copper
mg/kg
Lead
mg/kg
Magnesium (total)
g/kg
Manganese
mg/kg
Mercury
µg/kg
Nickel
mg/kg
Zinc
mg/kg
L
P
E

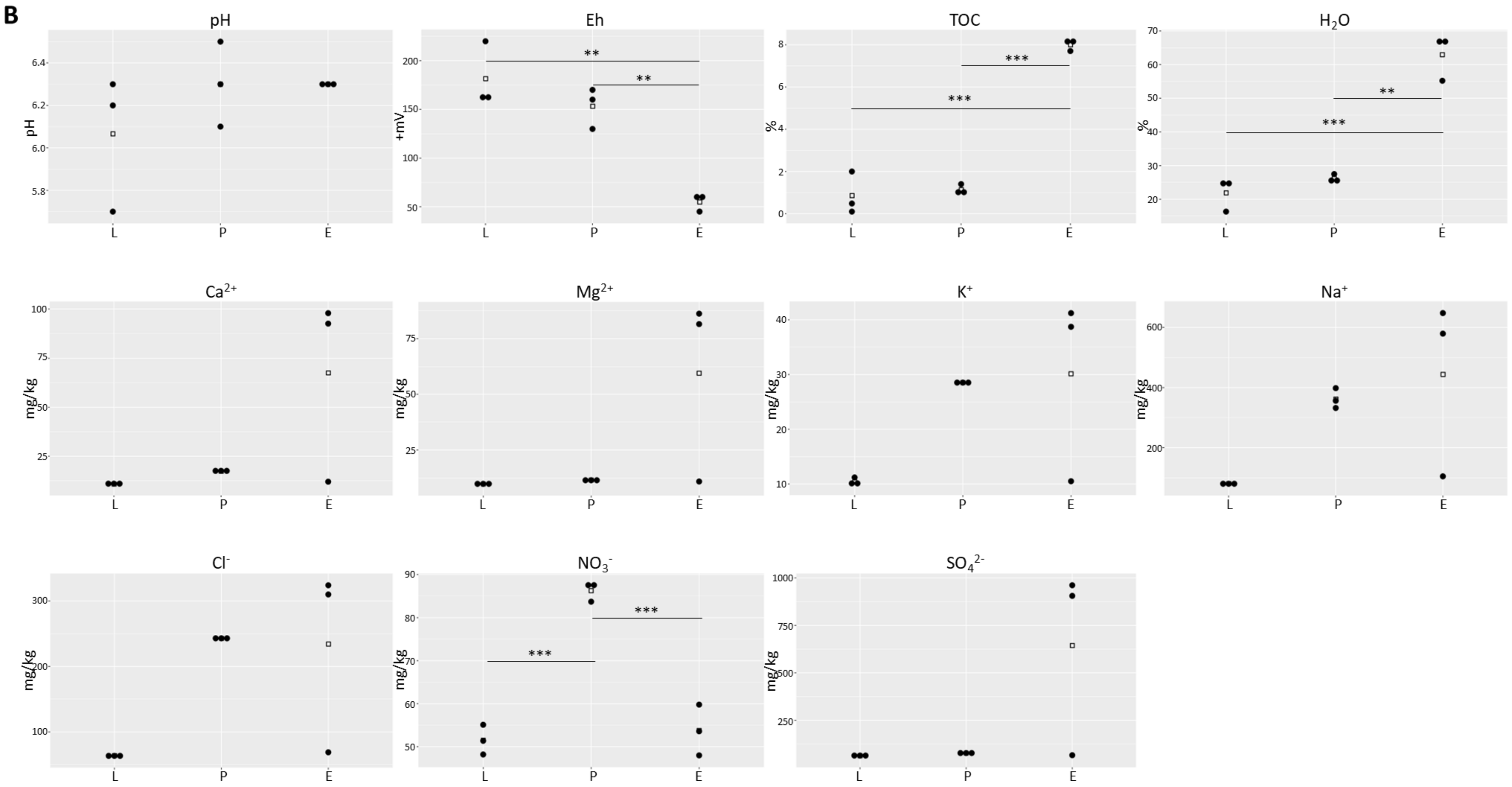


**Fig. S1: Physicochemical profile of sediment from sites ‘L’, ‘P’, and ‘E’.** (**A**) Metal concentrations measured in sediment samples. (**B**) Edaphic properties of pH, redox potential (Eh), total organic carbon (TOC), moisture content ($H_2O$), chloride, nitrate, sulfate, sodium, potassium, calcium, and magnesium. Panel A displays total magnesium whereas panel B displays cationic $Mg^{2+}$. Ammonium was not detected and is omitted. Individual replicates are displayed as ● (*n* = 3), means as □ (*p* values: ***<0.001, **<0.01, *<0.05, ANOVA for normal data with equal variance, and Kruskal-Wallis test for non-normal data).

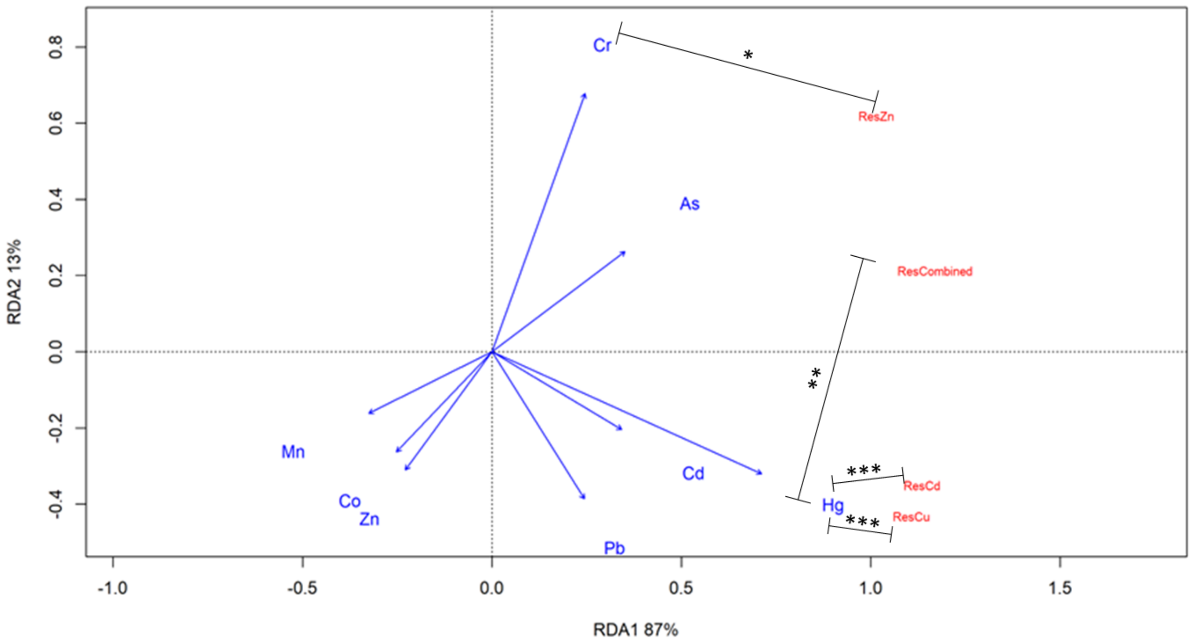


**Fig. S2: Positive correlation between metal-resistant colony forming unit and some heavy metal concentrations.** Redundancy analysis (RDA) was used to identify metals that best explain the variation in metal-resistant colony forming unit abundance. Only metals with the best explanatory value have been included for clarity. Significance values calculated from an independent Spearman's correlation coefficient *(p* values: ***<0.001, **<0.01, *<0.05, *n* = 3).

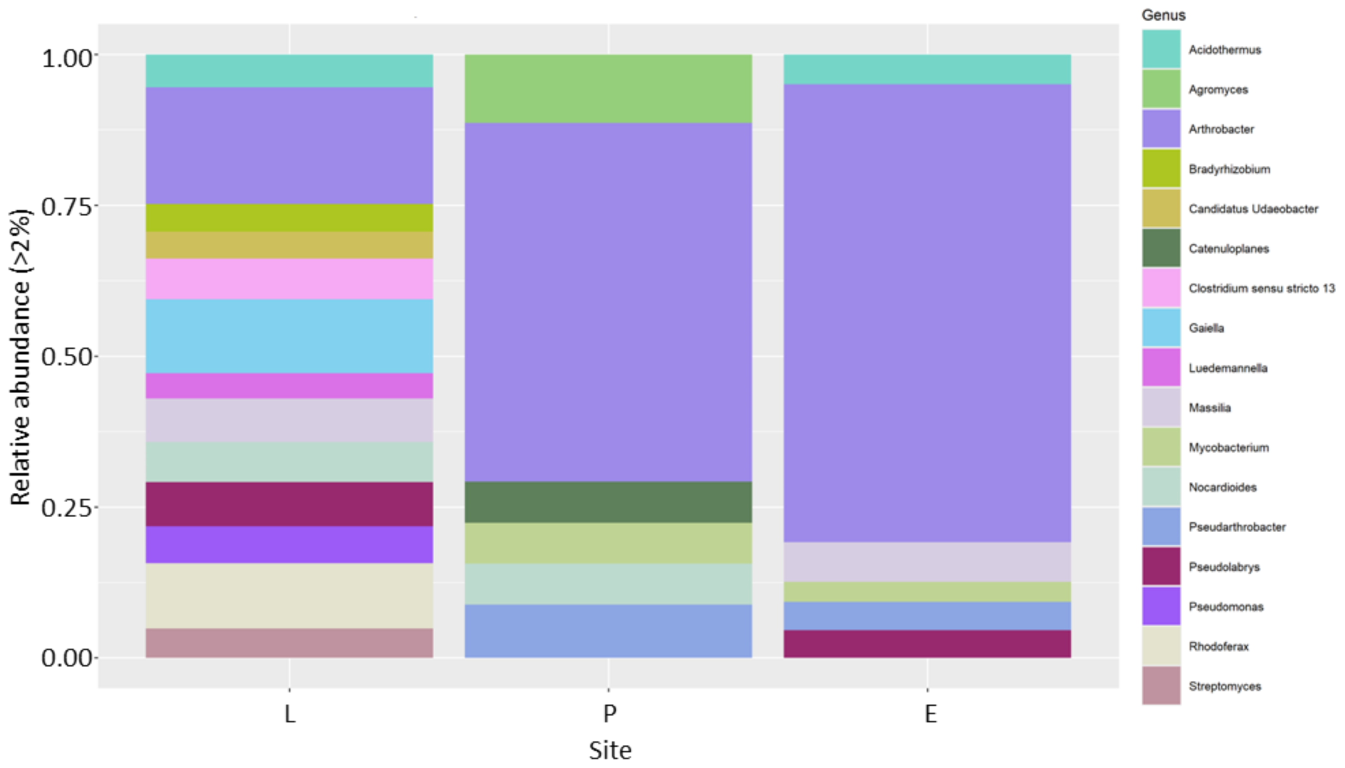


**Fig. S3: Reduced diversity of bacterial genera at industrial sites (P, E) based on metagenomic analysis.** Relative abundance of bacterial genera based on amplicon sequence variants from each site. Only genera with a relative abundance >2% are displayed and proportions are rescaled accordingly.

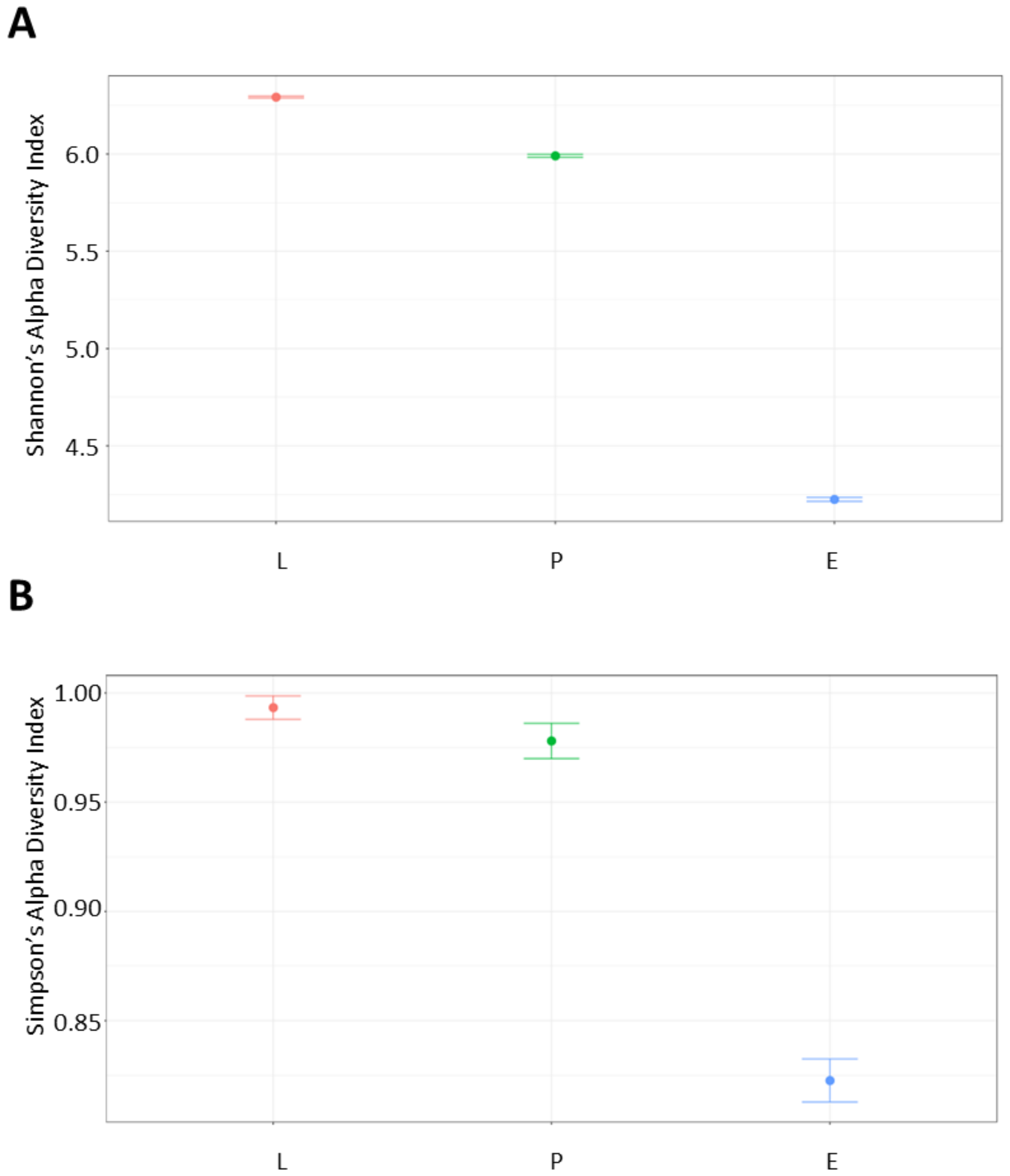


**Fig. S4: Alpha diversity of the bacterial communities in sediments from the industrial (P, E) and reference (L) sites.** Alpha diversity was assessed using (**A**) Shannon's (**B**) or Simpson's index with 95% confidence intervals (Hutcheson's t-test). All amplicon sequence variants that appeared more than two times were retained in this analysis.

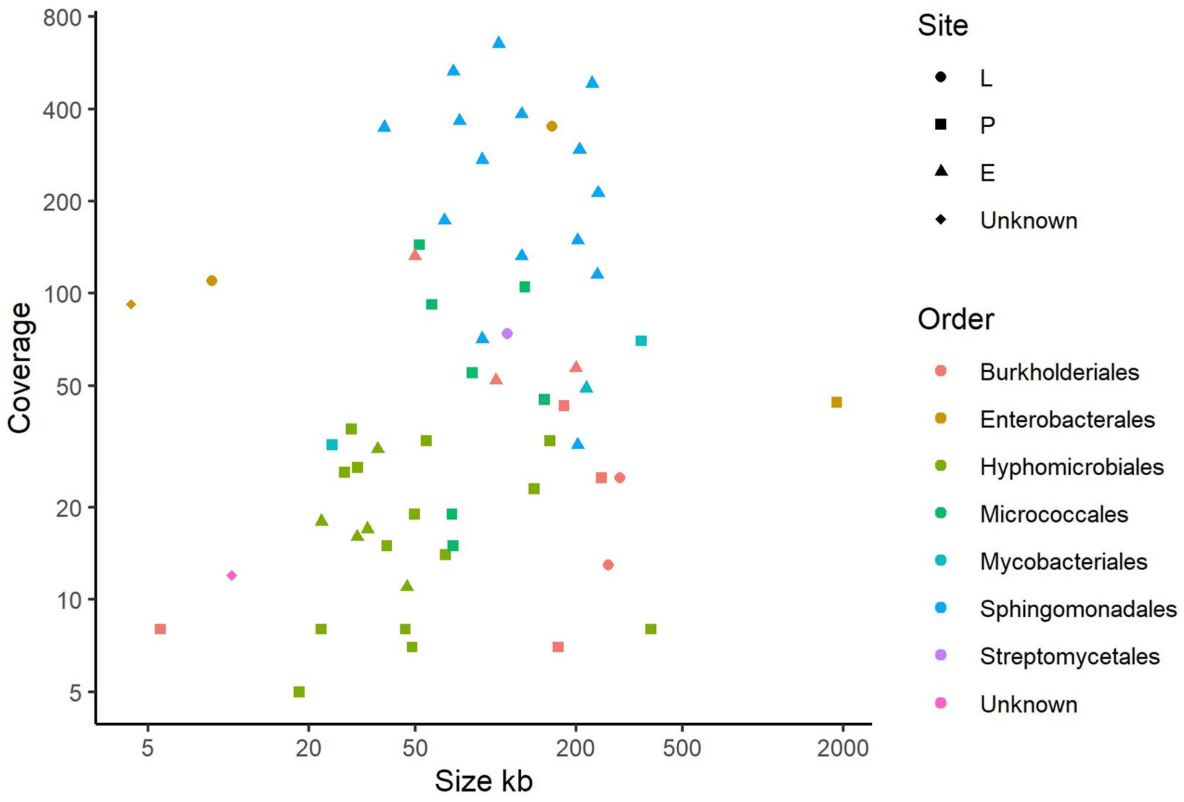


**Fig. S5: Plasmids were typically greater than 50 kb in size.** Plasmid sizes are plotted against Oxford Nanopore sequencing coverage using a double logarithmic scale. Plasmids are categorised by taxonomic order of their bacterial host (colour) and the sample site of origin (shape).

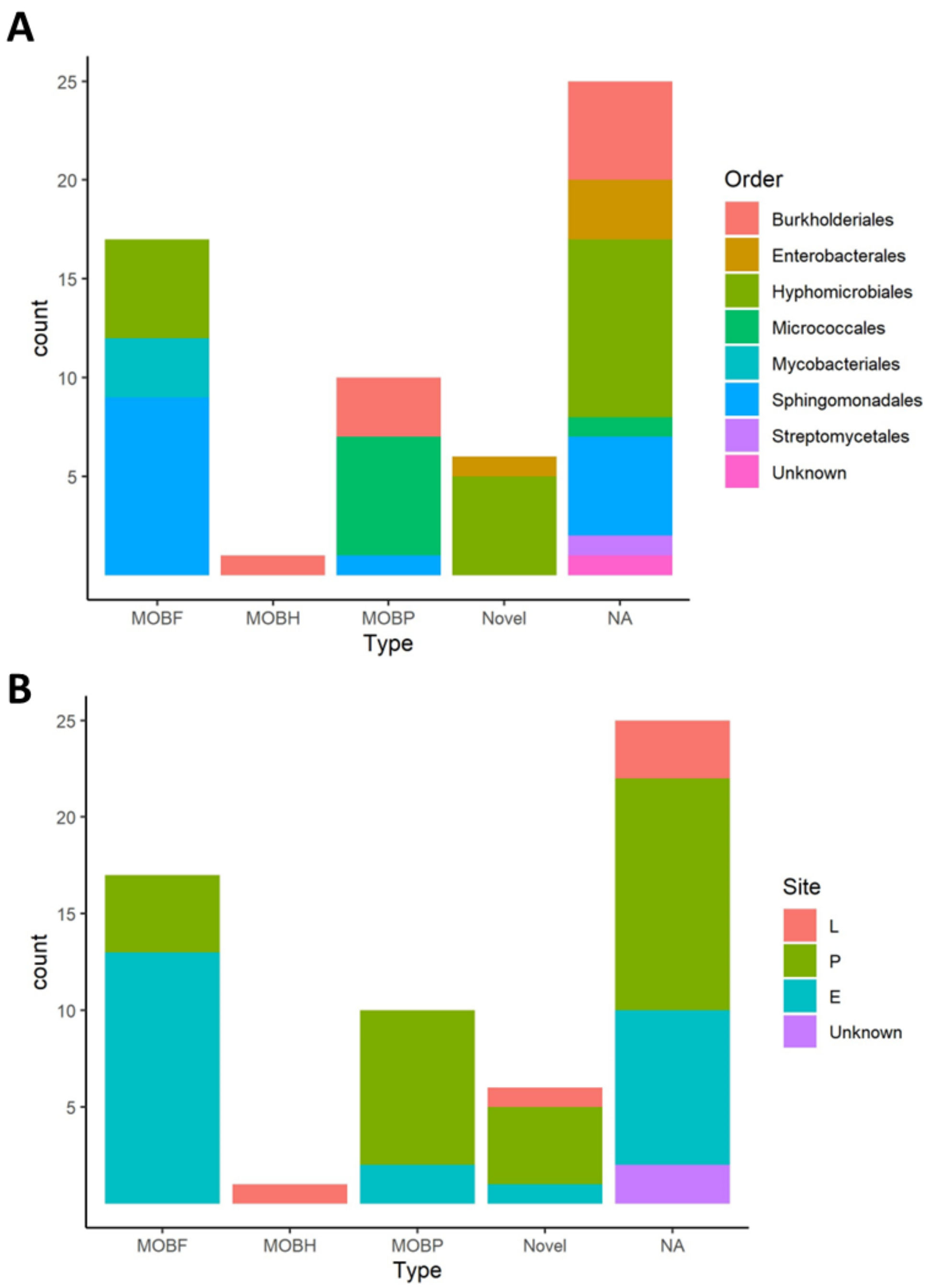


**Fig. S6: MOB$_F$ and MOB$_P$ plasmid types predominated among mobilisable plasmids.** Plasmids were grouped by mobility type predicted by MOBscan [41] and further subdivided by (**A**) bacterial host taxonomic order or (**B**) sample site. Plasmids without relaxase genes are classified as not applicable (NA), whereas plasmids with relaxase genes that did not belong to a known group are classified as novel.

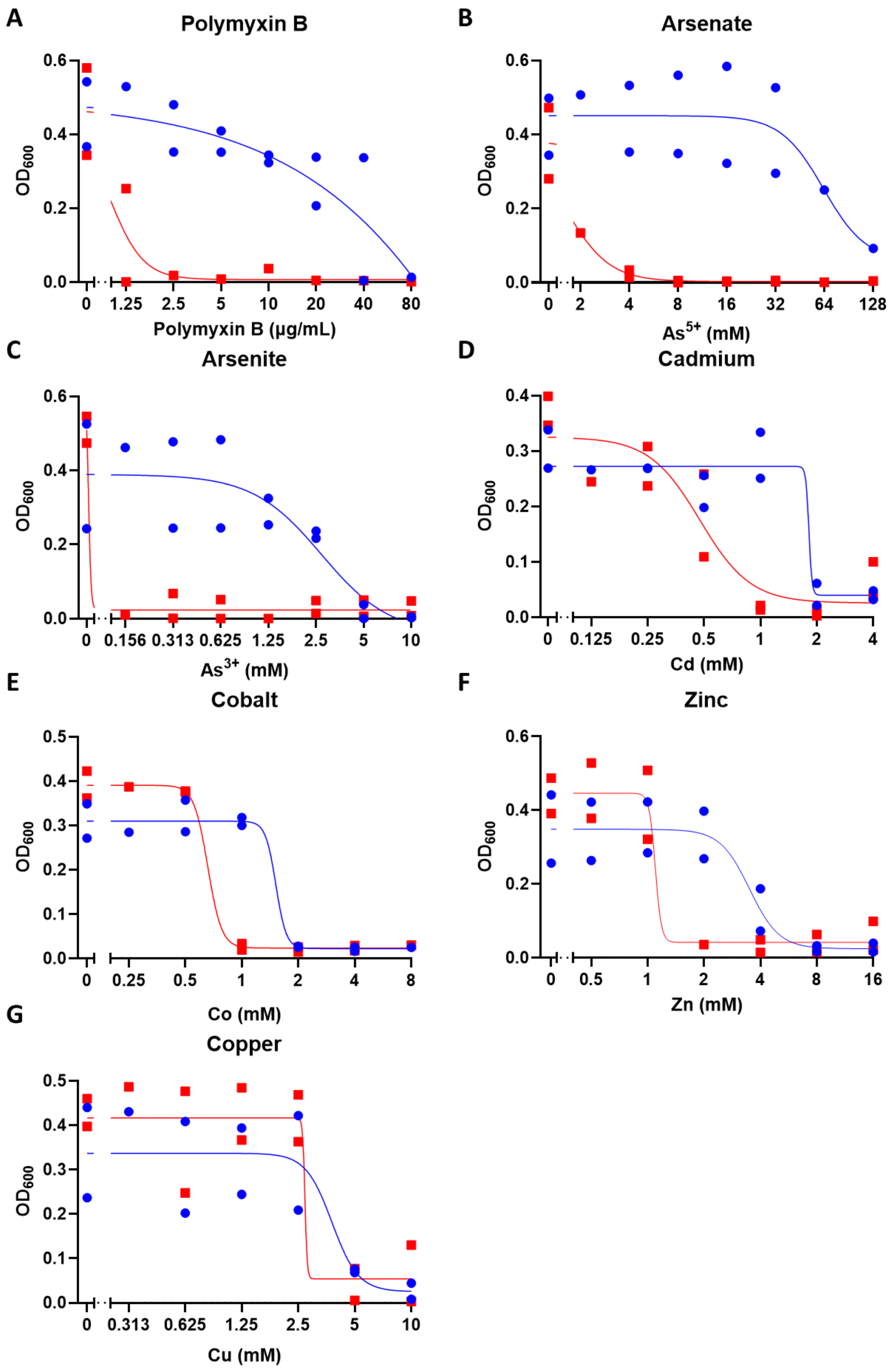


**Fig. S7: Plasmid pPLE30.2 is responsible for multi-metal and polymyxin resistance in *Pigmentiphaga litoralis* E30**. Cell growth measured at 48 hours for wild-type (●) and plasmid-cured strains (■) exposed to a concentration gradient of (**A**) polymyxin B, (**B**) arsenate, (**C**) arsenite, (**D**) cadmium, (**E**) cobalt, (**F**) zinc, and (**G**) copper. *n* = 2 biological replicates (mean of at least three technical replicates) plotted along with a sigmoidal non-linear regression.

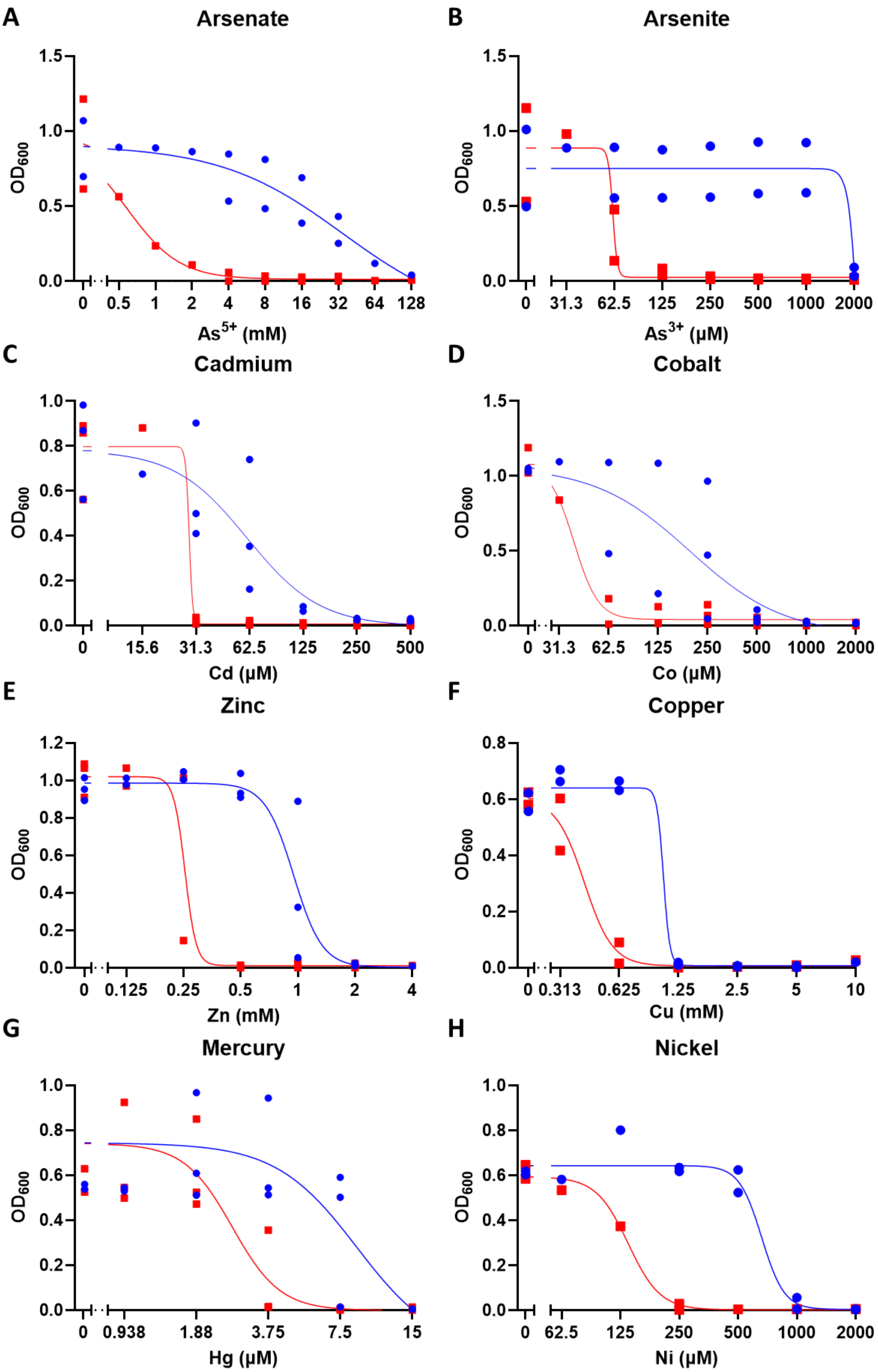


**Fig. S8: Plasmids are responsible for multi-metal resistance in *Sphingomonas hankookensis* E10.** Cell growth measured at 18 hours for wild-type (●) and plasmid-cured strains (■) exposed to a concentration gradient of (**A**) arsenate, (**B**) arsenite, (**C**) cadmium, (**D**) cobalt, (**E**) zinc, (**F**) copper, (**G**) mercury, and (**H**) nickel. $n$ = 2-3 biological replicates (mean of at least three technical replicates) plotted along with a sigmoidal non-linear regression.

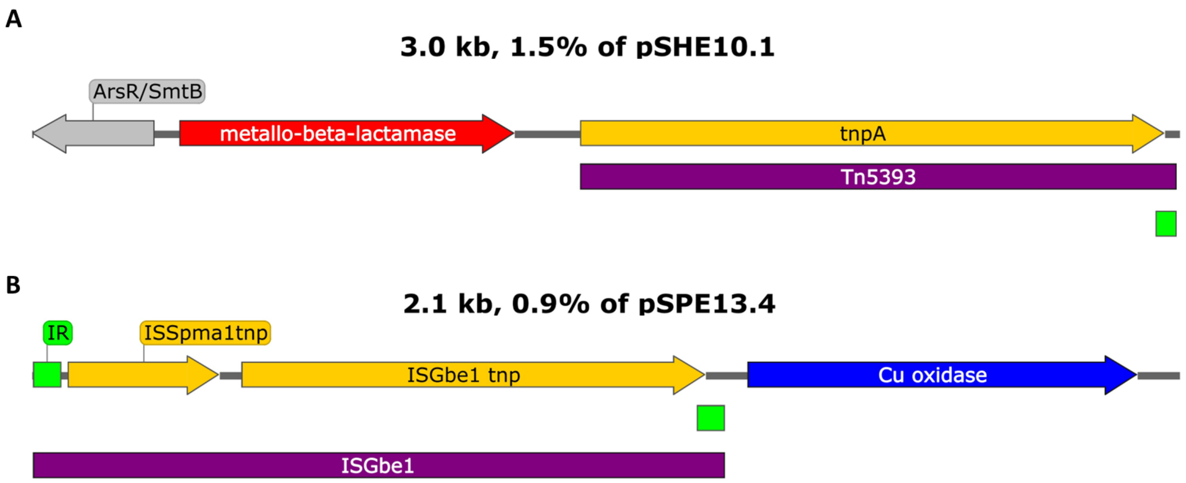


**Fig. S9: Resistance genes adjacent to insertion sequences and transposons.** Sections of sequence from (**A**) pSHE10.1 and (**B**) pSPE13.4. Arrows and blocks indicate bioinformatically predicted features, colour-coded by predicted function: purple blocks indicate predicted insertion sequences or transposons, orange arrows represent transposase genes, green blocks indicate inverted repeats, blue arrows represent metal resistance genes, red arrows represent antibiotic resistance genes, and grey arrows indicate unrelated genes.

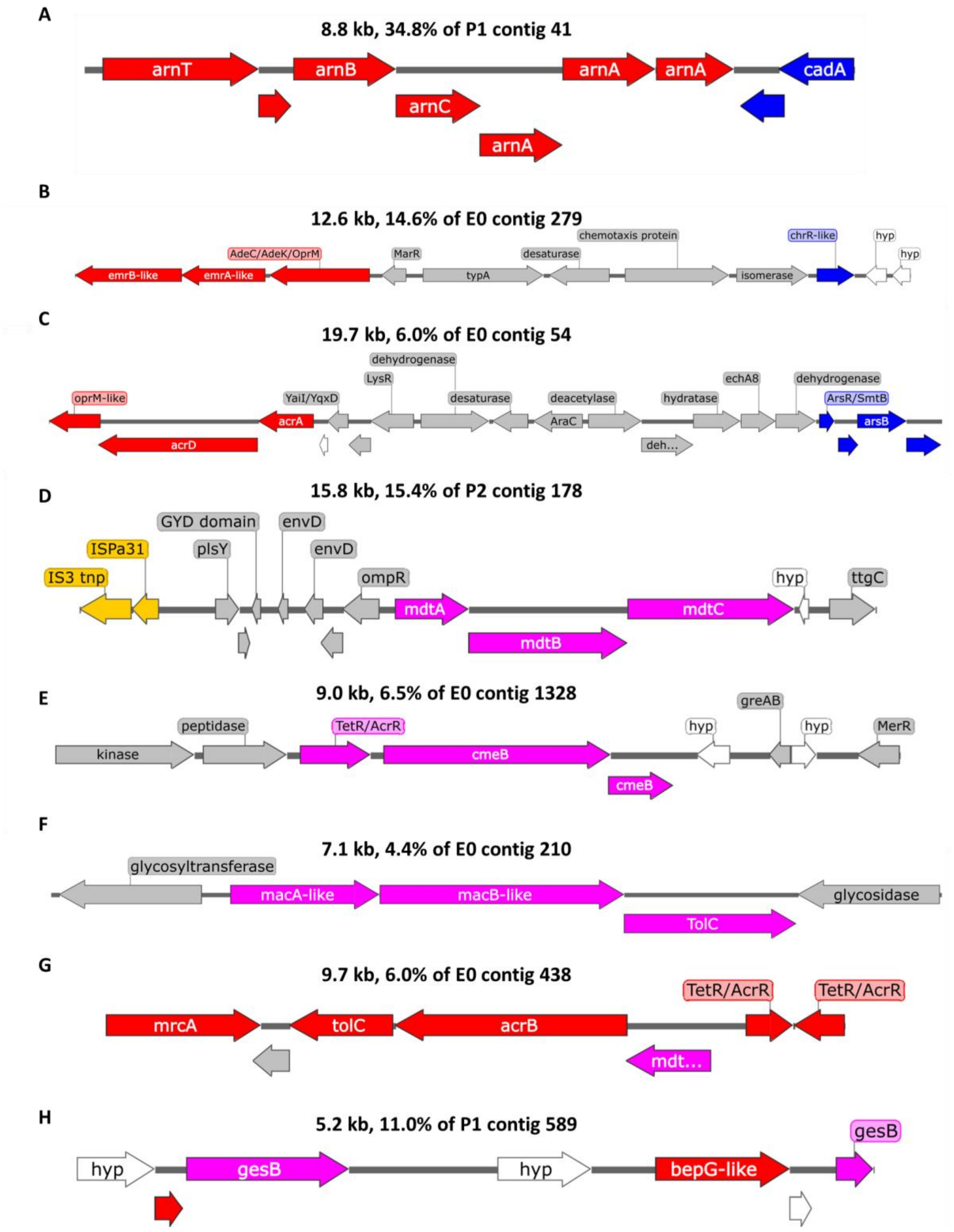


**Fig. S10: Resistance contigs exhibiting co-resistance (A-C, G-H), and cross-resistance (D-H) features**. Sequence sections are shown from (**A**) P1 contig 41, (**B**) E0 contig 279, (**C**) E0 contig 54, (**D**) P2 contig 178, (**E**) E0 contig 1328, (**F**) E0 contig 210, (**G**) E0 contig 438, and (**H**) P1 contig 589. Arrows indicate bioinformatically predicted genes, colour-coded by predicted function: pink represents antibiotic and metal cross-resistance genes, red indicates antibiotic resistance genes, blue represents metal resistance genes, orange indicates mobile genetic elements, grey represents unrelated genes, and white indicates genes of unknown identity.
